# SARS-CoV-2 ORF3a blocks lysosomal cholesterol egress by disrupting VPS39-regulated NPC2 trafficking and BMP metabolism

**DOI:** 10.1101/2024.11.13.623299

**Authors:** Baley A. Goodson, Valeria Montenegro Vazquez, Aliza Doyle, Oralia M. Kolaczkowski, Rui Liu, Jingyue Jia, Morié Ishida, Hu Wang, Xianlin Han, Chunyan Ye, Alison M. Kell, Steven B. Bradfute, Monica Rosas Lemus, Jing Pu

## Abstract

Cholesterol homeostasis relies on lysosomes, which release free cholesterol from degraded lipids. We show that SARS-CoV-2 blocks lysosomal cholesterol export through its protein ORF3a. ORF3a binds the HOPS subunit VPS39, and disrupting this interaction restores cholesterol trafficking. Two mechanisms underlie this defect. First, ORF3a–VPS39 interaction traps the sorting receptor CI-MPR and the retrieval complex retromer in endosomes/lysosomes, impairing trafficking of the cholesterol transporter NPC2. Retromer deletion reproduced these defects. Second, ORF3a reduces bis(monoacylglycerol)phosphates (BMPs), lysosomal lipids required for cholesterol export. Lipidomics and proteomics revealed altered metabolism of BMP precursors, mitochondrial phosphatidylglycerols (PGs), and reduced mitochondrial proteins at lysosomes. ORF3a–VPS39 interaction decreased lysosome–mitochondrion membrane contact sites (MCS), excluding autophagy or mitochondrion-derived vesicles as routes for PG transfer. VPS39 deletion decreased the MCS and BMPs. These findings identify VPS39 as a regulator of NPC2 trafficking and BMP biosynthesis and reveal that ORF3a disrupts both pathways to block cholesterol egress.

## INTRODUCTION

Cholesterol plays a vital role in maintaining cellular structure and functions. Thus, cells tightly regulate cholesterol levels and distribution through multiple pathways. Cholesterol pathways are not only disturbed in metabolic disorders but also hijacked in infectious diseases. COVID-19, caused by the SARS-CoV-2 virus, has been associated with significant disruptions in cholesterol metabolism^(1–5)^. In cases of long COVID, an increased risk of dyslipidemia and cardiovascular diseases is marked by elevated low-density lipoprotein (LDL) cholesterol and reduced high-density lipoprotein (HDL) cholesterol levels^(4,5)^.

Lysosomes, the degradation organelles, receive and break down internalized lipoprotein particles and cellular macrolipids delivered via endocytosis and autophagy, respectively, releasing free cholesterol in the process^(6)^. The released cholesterol is then distributed to other cellular compartments to support structural and metabolic demands^(7,8)^. The cholesterol delivered to the endoplasmic reticulum (ER) serves as a signal to regulate the expression of genes that control cholesterol synthesis and uptake^(9)^. The cholesterol supplied to the plasma membrane is the major source for maintaining the local cholesterol pool that modulate plasma membrane fluidity and stability^(10)^. SARS-CoV-2 exploits plasma membrane cholesterol to enhance infectivity^(11)^, as evidenced by a reduced binding between the viral spike protein and its host receptor after plasma membrane cholesterol depletion^(12)^. Therefore, the lysosomal cholesterol transport is central to cellular cholesterol homeostasis and may also play a role in SARS-CoV-2 infection.

Lysosomal cholesterol egress is the first step of cholesterol distribution. This process is primarily mediated by the lysosomal transmembrane cholesterol transporter Niemann-Pick C1 (NPC1) and the lumenal cholesterol-binding protein Niemann-Pick C2 (NPC2), which act cooperatively to transfer cholesterol out of lysosomes^(8)^. Additional lysosomal proteins, including the lumenal saposins and membrane proteins LAMP2^(13,14)^, also contribute to cholesterol egress^(8)^. The proper trafficking and localization of newly synthesized lysosome transmembrane proteins depend on sorting signals and clathrin coats^(15)^. In contrast, the lysosome lumenal proteins are directed by sorting receptors, such as the mannose-6-phosphate receptors (M6PRs)^(15)^. These receptors deliver lysosome lumenal proteins from the trans-Golgi network (TGN) to endosomes and are subsequently recycled back to the TGN. This recycling requires retromer complexes at endosomes and lysosomes^(16–19)^ and Golgi-associated retrograde protein (GARP) complex at the TGN^(20)^. Disruption of these transport pathways by impairing M6PR, HOPS or GARP functions compromises the delivery of newly synthesized proteins to lysosomes and results in accumulation of cholesterol^(18,21,22)^. Such lysosomal cholesterol sequestration profoundly disrupts lipid homeostasis and impairs cell functions, a defining feature of Niemann-Pick disease type C, caused by mutations in NPC1 or NPC2^(23,24)^. Beyond genetic disorders, certain viruses exploit lysosomal cholesterol transport pathways to complete their life cycles. For instance, NPC1 is hijacked by Ebola virus for their infection^(25)^.

In addition to protein-mediated cholesterol transport, the phospholipid bis(monoacylglycero)phosphates (BMPs, also called lysobisphosphatidic acids or LBPAs), play an important role in lysosomal cholesterol egress. BMPs are highly enriched in the late endosome and lysosome membrane, where they contribute to lysosome sorting and degradation functions^(9,26–28)^. BMP has also been implicated in viral infection by promoting viral fusion with late endosome membranes^(29–32)^. Beyond these function, BMPs directly interact with NPC2 to promote cholesterol egress in an NPC1-independent way, as supplementation with BMPs rescues cholesterol sequestration in NPC1-defecient cells but has no effect in NPC2-defecient cells^(33,34)^. The synthesis and turnover of BMPs depend on multiple lysosomal enzymes that convert the mitochondrial phospholipids phosphatidylglycerols (PGs) into BMPs and degrade BMPs through hydrolysis^(35–39)^.

In this study, we show that SARS-CoV-2 infection drives cholesterol sequestration in lysosomes, primarily through the viral protein ORF3a. ORF3a binds the HOPS subunit VPS39, disrupting lysosomal lumenal protein trafficking by retaining the cation-independent mannose 6-phosphate receptor (CI-MPR) and retromer complex at endosomes and lysosomes. In addition, ORF3a–VPS39 interaction impairs lysosome-mitochondrion membrane contact sites, potentially limiting lipid transport from mitochondria to lysosomes required for BMP synthesis. By simultaneously disturbing these two VPS39-dependent pathways, ORF3a effectively sequesters cholesterol within lysosomes, revealing a mechanistic link between SARS-CoV-2 infection and lysosomal lipid dysregulation.

## RESULTS

### Increased lysosomal cholesterol in SARS-CoV-2-infected cells

To examine how SARS-CoV-2 infection alters host cholesterol at cellular levels, we infected human lung carcinoma epithelial cells A549, stably expressing the viral entry receptor ACE2 of human species (A549-hACE2), with the Washington strain of SARS-CoV-2. We confirmed infection in approximately 80% of cells by immunostaining for double-stranded RNA (dsRNA) (Figs. 1A & S1A). Using filipin staining to detect free cholesterol, we observed a marked increase in filipin signal in infected cells, appearing as punctate structures, compared to mock-treated cells (Fig. 1A). Similar observations were made in a different ACE2-overexpressing A549 cell line (Fig. S1B). Co-staining with an antibody against late endosomal and lysosomal membrane protein LAMP1 revealed that the filipin puncta colocalized with LAMP1-positive vesicles (Fig. 1A), indicating that cholesterol was sequestrated in the late endosomes and lysosomes. Due to the similarity of these two organelles, we refer them to lysosomes hereafter. Quantification of confocal images showed that lysosomal cholesterol levels significantly increased at 24– and 48-hours post-infection (Fig. 1B). Although filipin puncta were also present at 72 hours post-infection, mock cells showed elevated filipin signal, likely due to cell responses associated with high cell density^(40)^ (Fig. 1B). Further analysis of over 1,500 infected cells using a high-content imaging system confirmed a consistent increase in filipin-positive puncta (Fig. 1C). To validate this cholesterol alteration, we infected monkey kidney cells Vero E6 with SARS-CoV-2, resulting approximately 80% infection rate (Fig. S1C). High-content imaging revealed that the infected cells displayed a significant 50–60% increase in cholesterol relative to LAMP1, compared with mock-treated cells, at 18 and 24 hours post-infection. (Figs. 1D, 1E, S1D), confirming that SARS-CoV-2 infection induces an increase in lysosomal cholesterol. This change persisted beyond the viral eclipse phase (∼10 hours)^(41)^. Gas chromatography-mass spectrometry analysis of total cholesterol levels in mock and infected A549 cells showed no significant difference (Fig. 1F), suggesting that SARS-CoV-2 infection mainly alters cholesterol distribution within cells, possibly due to impaired lysosomal cholesterol transport.

**Figure 1.**
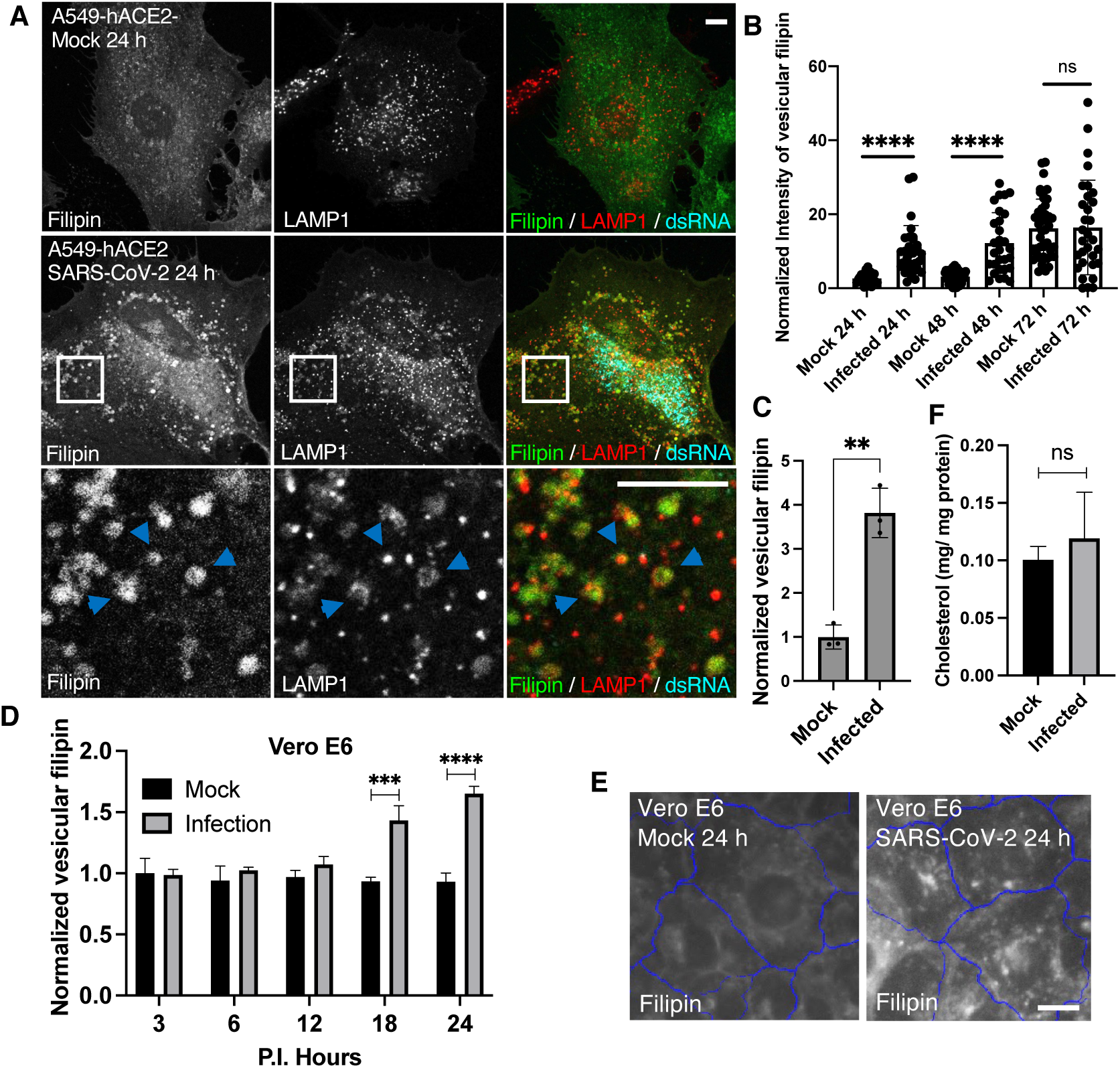
Characterization of cellular free cholesterol in SARS-CoV-2 infected cells. **A-C.** A549 cells stably expressing human ACE2 receptor (A549-hACE2) were infected with SARS-CoV-2 and fixed at the indicated time post-infection (P.I.). Filipin was applied to visualize free cholesterol. A dsRNA antibody was used to identify the infected cells. A LAMP1 antibody was used to label lysosomes. Confocal microscopy was performed to collect fluorescence images, and the images at 24 h P.I. were shown in **A**, with arrows pointing colocalized filipin and LAMP1. The intensity of Filipin puncta was quantified with FIJI, and the mean values of total filipin in each cell from three independent experiments were plotted, shown as in **B**. Vesicular filipin was quantified at 24 h P.I. from at least 1,000 infected cells per well and three replicate wells in an experiment by CellInsight microscope platform. Three independent experiments were performed, and pooled results were shown in **C**. **D,E.** Similar experiments were perform as above in Vero E6 cells, and filipin was quantified at the indicated time points. **F.** Total lipids were extracted from mock and infected A549-hACE2 cells at 24 h P.I. and subjected to free cholesterol measurement with GC-MS. Total protein levels were measure by BCA assays. Two independent experiments with 3 replications were performed. Bar graphs are presented as min-max (**B**) or mean ± SD (others). *p* values were determined using *t’* test or One-way ANOVA test. **, *p*<0.001, ***, *p*<0.001, ****, *p*<0.0001, n.s., not significant. Scale bars, 5 μm.

### Viral protein ORF3a is responsible for increased lysosomal cholesterol

To identify the specific viral proteins responsible for lysosomal cholesterol sequestration, we transfected A549-hACE2 cells with 28 plasmids individually, which encode SARS-CoV-2 proteins, and quantified filipin puncta 24 hours post-transfection. Most viral proteins (24 out of 28) did not show a significant effect on cholesterol distribution. NSP10 and NSP13 had modest effects, ORF10 doubled the filipin signal, and ORF3a produced the strongest increase in filipin puncta (Fig. 2A). ORF3a, expressed from a SARS-CoV-2 subgenomic RNA, is an accessory protein. Unlike NSP10, NSP13, or ORF10, which were predominantly cytoplasmic (Fig. S2), ORF3a displayed punctate or halo-like structures that partially co-localized with LAMP2 (Fig. 2B), consistent with previous reports of ORF3a localization to endosomes/lysosomes^(42–44)^. Remarkably, ORF3a-positive lysosomes appeared swollen and were filled with filipin (Fig. 1B, arrow in enlarged frame 1), in contrast to ORF3a-negative lysosomes in the same cells, which remained small and displayed minimal filipin signal (Fig. 1B, arrow in enlarged frame 2). This suggests that ORF3a, particularly when localized on lysosomes, plays a critical role in cholesterol sequestration.

**Figure 2.**
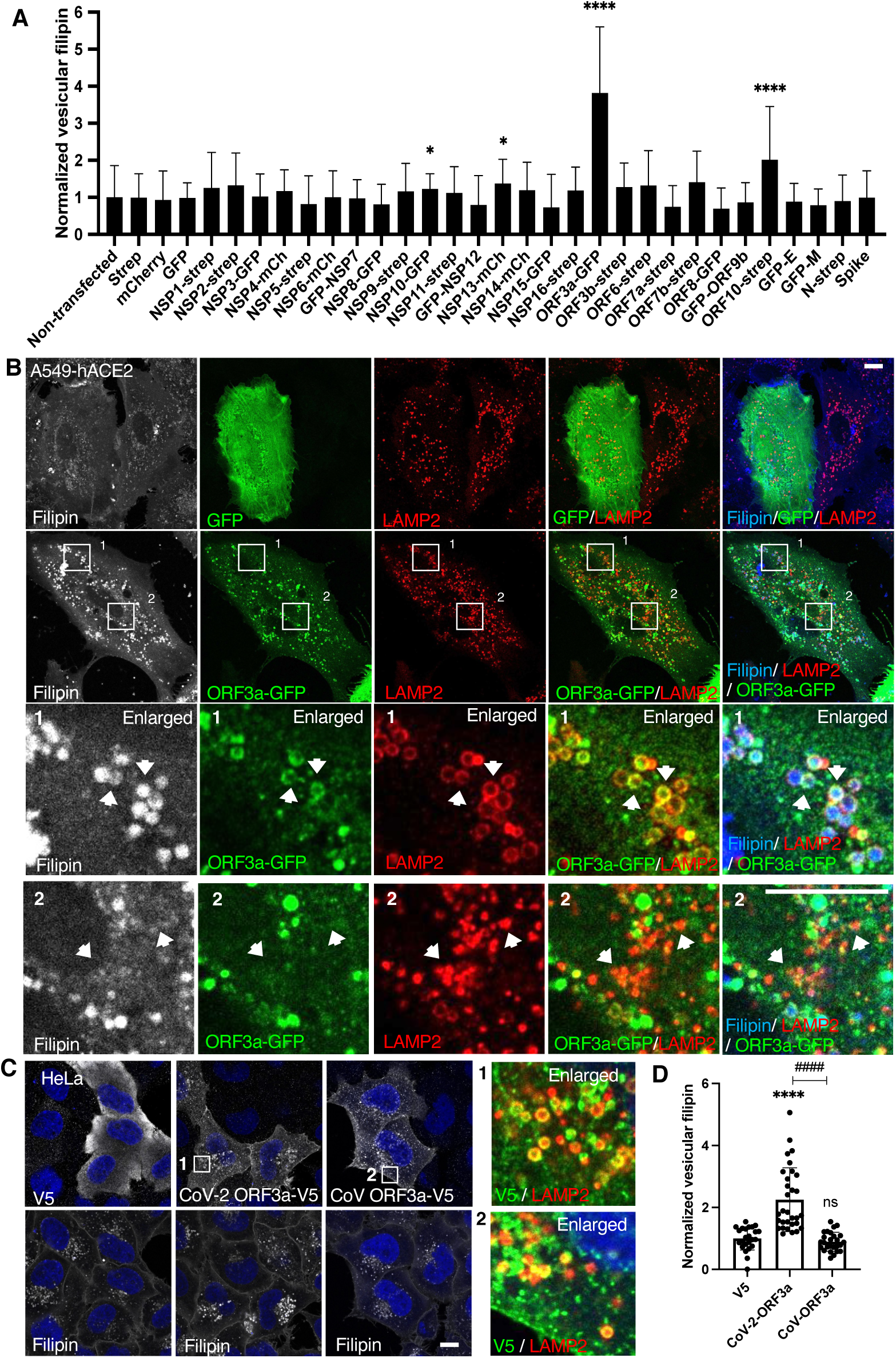
Identification of SARS-CoV-2 proteins responsible for lysosome cholesterol sequestration. **A.** Plasmids encoding SARS-CoV-2 proteins were transfected into A549-hACE2 cells individually. Cells were fixed at 24 h post-transfection and co-stained with filipin to visualized free cholesterol and antibodies against the epitope tags to identify transfected cells. Spike protein has no tag and was identified with an antibody. Confocal imaging was performed to randomly take images, and filipin intensity was quantified with FIJI in twenty to fifty cells from two independent experiments. **B.** A549-hACE2 cells were transfected with ORF3a-GFP plasmid as in **A** and co-stained with filipin and GFP and LAMP2 antibodies. **C,D**. HeLa cells were transfected with plasmids encode V5 fused ORF3a proteins in SARS-CoV or SARS-CoV-2. V5 alone served as a control. Staining and quantification was performed as described above. Bar graphs are presented as mean ± SD. *p* values were determined using *t’* test or One-way ANOVA test. *, *p*<0.05, **** (vs. control) or #### (CoV vs CoV-2), *p*<0.0001, n.s., not significant. Scale bars, 5 μm. Framed images in C were enlarged 9.36 folds.

To further validate this observation, we transfected HeLa cells with an ORF3a plasmid tagged with a V5 epitope. ORF3a-V5 also formed punctate or halo structures, partially co-localized with LAMP2, and significantly increased filipin signal (Fig. 2C,D), demonstrating that ORF3a enhances lysosomal cholesterol accumulation across different cell types. Additionally, we included ORF3a from SARS-CoV, which shares over 70% sequence identity with SARS-CoV-2 ORF3a, as a comparison. Interestingly, while SARS-CoV ORF3a also displayed a punctate distribution, it did not increase filipin (Fig. 2C,D), suggesting a specificity in cholesterol sequestration by SARS-CoV-2 ORF3a.

Together, these results demonstrate that SARS-CoV-2 infection sequesters host lysosomal cholesterol primarily through the action of ORF3a. Expression of ORF3a alone is sufficient to drive cholesterol accumulation in lysosomes, likely due to its lysosomal localization. The differential effects on lysosomal cholesterol between ORF3a proteins from SARS-CoV and SARS-CoV-2 may offer insights into the underlying mechanisms.

### The Structural difference between SARS-CoV and SARS-CoV-2 ORF3a

Previous studies revealed that SARS-CoV-2 ORF3a, but not SARS-CoV ORF3a, interacts with the host protein VPS39^(42–45)^ (Figs. 3A & S3B), a subunit of the HOPS complex. Our recent work showed that the HOPS complex is essential for lysosomal cholesterol egress, using VPS39 CRISPR-knockout (KO) cells^(18)^. SARS-CoV-2 infection did not downregulate the expression of VPS39 or VPS11, another HOPS subunit (Fig. S3A). These results suggest that SARS-CoV-2 ORF3a disrupts lysosomal cholesterol egress by inhibiting VPS39 functions. To test this, we aimed to disrupt the ORF3a-VPS39 interaction by introducing specific point mutations.

**Figure 3.**
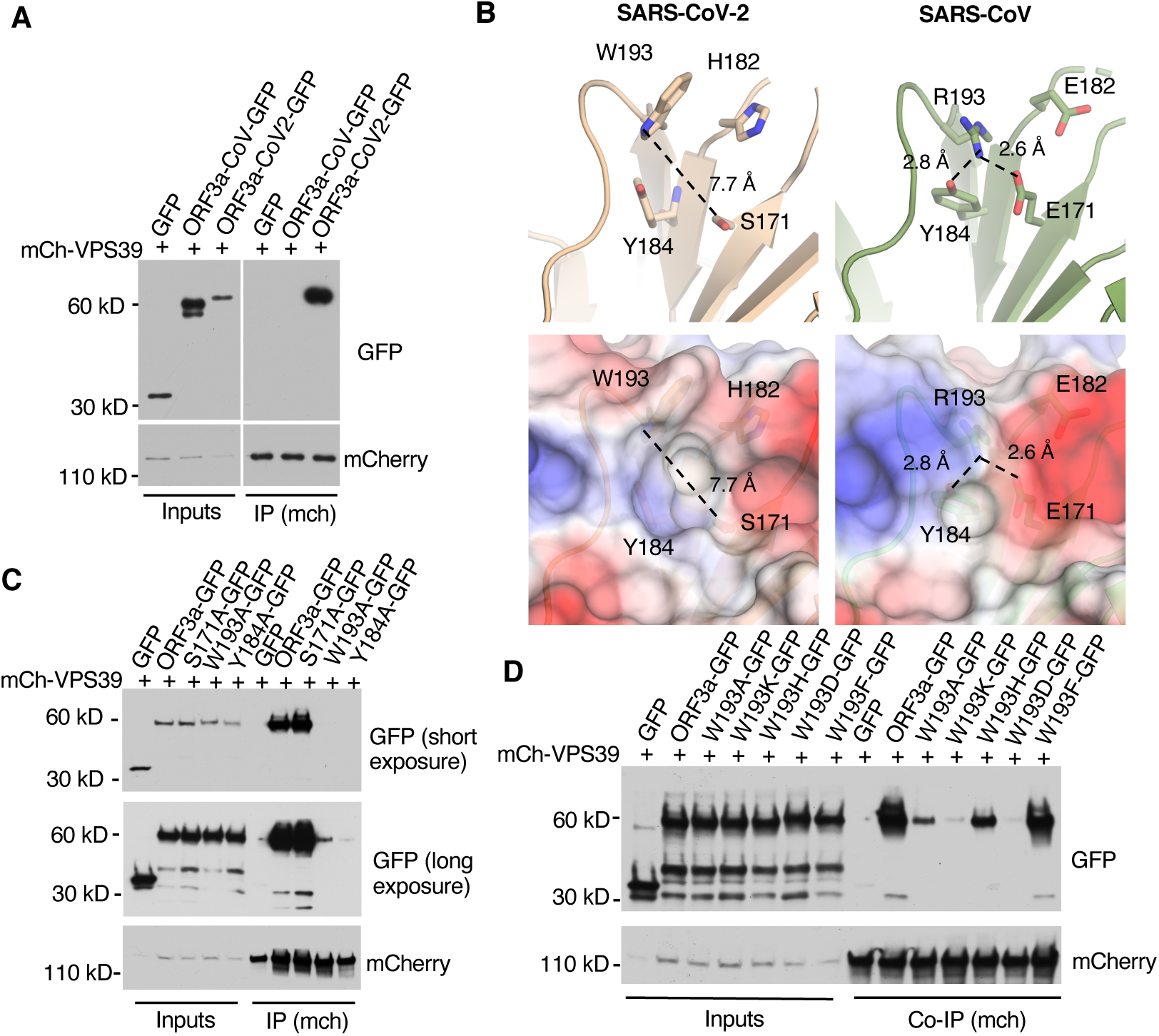
Structural analysis of ORF3a-VPS39 interaction interface in ORF3a. **A.** HeLa cells were co-transfected with mCh-VPS39 and different GFP-ORF3a constructs as indicated and lysed in a non-denature detergent-contained buffer for co-immunoprecipitation of mCh-VPS39. Immunoblotting was performed to detect GFP signals. **B.** The structures of ORF3a proteins from SARS-CoV-2 (beige) and SARS-CoV (green) were shown with the cartoon representation of the indicated residues (upper panel). The side chains are depicted as sticks with color code for carbon (beige or green), nitrogen (blue), and oxygen (red). The presence of W193 in SARS-CoV-2 induces an open conformation of the divergent loop that is 7.7 Å away from S171 (dotted line), while R193 in SARS-CoV forms hydrogen bonds with D171 or Y184 (dotted lines). The lower panels represent overlays between electrostatics surface and the cartoon representations. Positively charged surface is depicted in blue, hydrophobic surface in white, and negatively charged surface in red. All images and measurements were performed using PyMol V 2.4.0. **C,D.** Transfection and co-immunoprecipitation was performed as described in A to detect the interaction between mCh-VPS39 and GFP-tagged ORF3a and its mutants.

Miao et al. identified the residues S171 and W193 in SARS-CoV-2 ORF3a as crucial for its interaction with VPS39, as mutating these residues to those in SARS-CoV ORF3a abolished the interaction^(43)^. Structural analysis of SARS-CoV-2 ORF3a^(46,47)^ revealed that W193, Y184, S171, and H182 cluster near each other (Fig. 3B, upper), with W193 and Y184 forming a hydrophobic surface likely involved in protein-protein interactions, while S171 and H182 are positioned outside this hydrophobic surface (Fig. 3B, lower). In our alanine mutation screen, we found that W193A and Y184A mutations weakened or disrupted the interaction, whereas S171A or H182A mutations did not (Figs. 3C & S3C,D). This indicates that W193 and Y184, but not S171 or H182, are essential for binding. Further structural analysis indicates that the hydrophobic surface formed by W193 and Y184 (7.7 Å) in SARS-CoV-2 ORF3a is occluded in SARS-CoV ORF3a due to a hydrogen bond between R193 and E171 (2.6 Å) and Y184 (2.8 Å), resulting a “closed conformation” (Fig. 3B, upper). This model is supported by the observation that the S171E mutation impairs the interaction^(43)^, likely due to the large hydrophilic side chain of glutamic acid obstructing the hydrophobic surface formed by W193 and Y184, thereby blocking the interaction. Collectively, these findings suggest that W193 and Y184 form the primary binding site, while S171 and H182 contribute to an optimal binding environment.

Given that both W193 and Y184 are aromatic amino acids, we hypothesized that their aromatic rings play a role in the interaction. To test this, we mutated W193 to various amino acids and evaluated the effects on binding. Replacing W193 with another hydrophobic aromatic amino acid, such as phenylalanine, preserved the interaction. However, substituting W193 with a charged aromatic amino acid, like histidine, or with a hydrophobic but non-aromatic amino acid, like alanine, reduced binding. Furthermore, replacing W193 with a charged, non-aromatic amino acid, such as lysine or aspartic acid, nearly abolished the interaction (Fig. 3D). These results suggest that both the aromatic ring and hydrophobicity of W193 are critical for the binding of SARS-CoV-2 ORF3a to VPS39.

### ORF3a-VPS39 interaction interrupts NPC2 trafficking

With the understanding of structural basis of the interaction interface, we chose W193A mutation to block the interaction between ORF3a and VPS39. To minimize potential artifacts from overexpression, we individually introduced SARS-CoV ORF3a, SARS-CoV-2 ORF3a, the SARS-CoV-2 ORF3a W193A mutant, and a linker peptide that serves as a control into HeLa-Flp-In cells to generate tetracycline-inducible protein-expression cell lines. This system controls protein expression levels via a pre-engineered recombination site in the cells^(48,49)^. Likely due to low protein expression, we could not detect the ORF3a proteins by immunofluorescence (Fig. S4A), but immunoblotting showed expected bands in response to induction by the tetracycline analog doxycycline (Fig. 4A). Using immunoprecipitation, we confirmed that, in the Flp-In cells, CoV-2 ORF3a interacted with endogenous VPS39, and the W193A mutation abolished this interaction (Fig. 4B). High-content imaging quantification revealed that doxycycline-induce expression of ORF3a increased filipin signaling by 50-60% compared to the control cells (Figs. 4C,D & S4B). Importantly, the W193A mutant significantly reduced ORF3a-induced cholesterol accumulation (Figs. 4C,D & S4B), demonstrating that the ORF3a-VPS39 interaction is responsible for the cholesterol defect.

**Figure 4.**
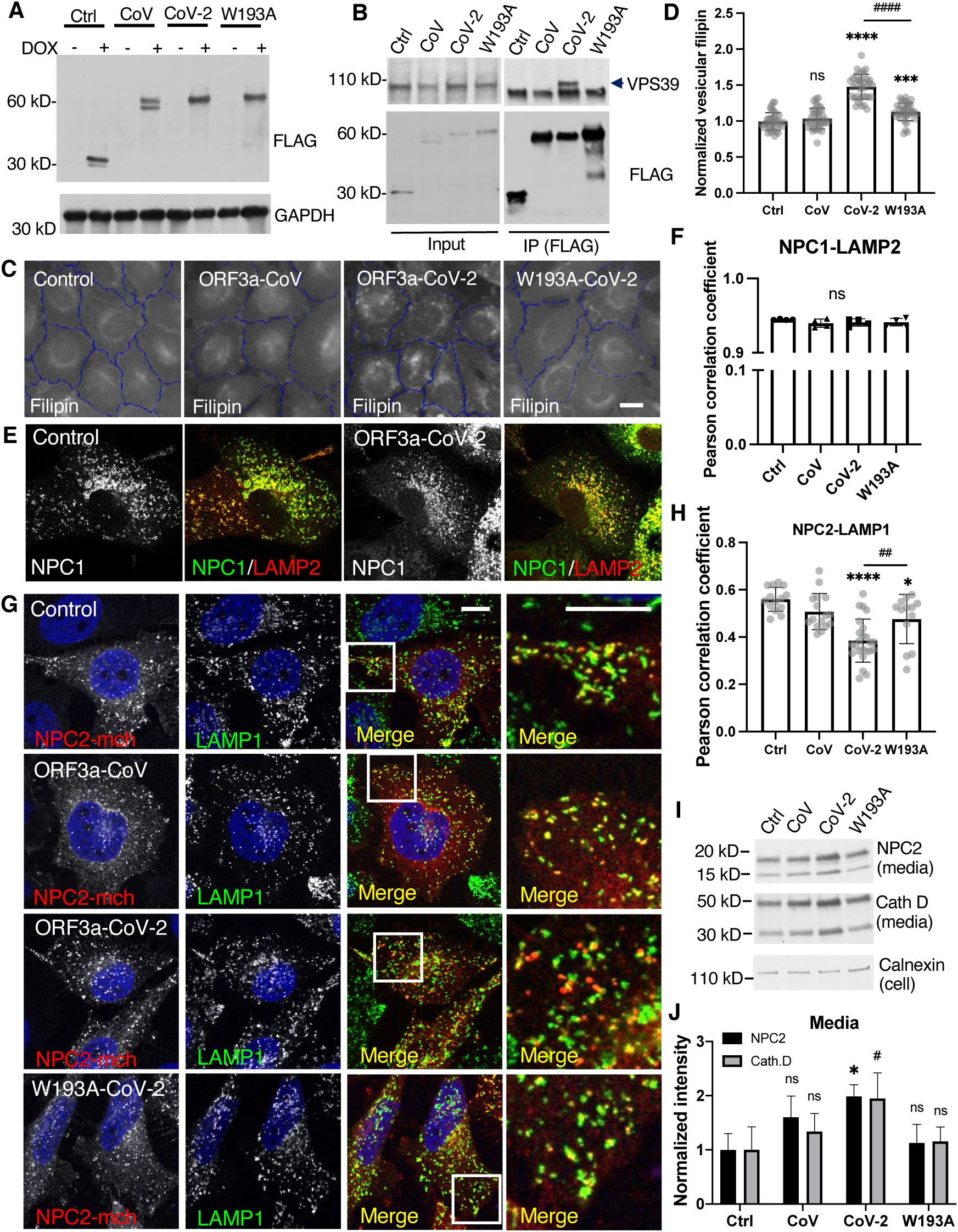
Characterization of NPC proteins in ORF3a-inducible expression cells. **A.** HeLa-Flp-In cells were established as described in Methods and treated with or without 1 µg/ml doxycycline for 16 hours. Immunoblotting was performed with the indicated antibodies. **B.** Co-immunoprecipitation was performed with FLAG antibody (M2)-coated magnetic beads. Endogenous VPS39 was detected by immunoblotting (arrow). **C,D.** HeLa-Flp-In cells were fixed and stained with filipin, CellMask, and DAPI for high-content imaging. Vesicular filipin was quantified in at least 1,000 cells per well, 6 wells per experiments, 5 independent experiments. **E,F.** Antibodies against NPC1 or LAMP2 were used for immunostaining in fixed cells. Confocal microscopy (**E**) and high-content imaging (**F**) was performed to analyze colocalization between NPC1 and LAMP2. **G,H.** Cells were transfected with NPC2-mCherry plasmid and immunostained with antibodies of mCherry and LAMP1. Confocal microscopy was performed, and FIJI was used to quantify colocalization between NPC2-mCherry and LAMP1 in the cells from two independent experiments. **I,J.** Cell culture media was collected and subjected to immunoblotting using the antibodies indicated. FIJI was used to quantify the bands from 3 replications in each of three independent experiments. Bar graphs are presented as mean ± SD. *p* values were determined using *t’* test (D,H, CoV-2 vs W193A) or One-way ANOVA test. * or #, *p*<0.05, ** or ##, *p*<0.01, \*\*\**, p*<0.001, **** (vs. control) or #### (CoV-2 vs W193A), *p*<0.0001, n.s., not significant. Scale bars, 5 μm. Control (Ctrl), linker peptide, CoV, SARS-CoV ORF3a, CoV-2, SARS-CoV-2 ORF3a, W193A, SARS-CoV-2 ORF3a-W193A mutant.

Previous research has shown that ORF3a expression increases lysosome pH^(42)^ and compromises lysosomal membrane integrity^(50)^, raising the question of whether lysosomal membrane damage or dysfunction leads to cholesterol sequestration. To investigate this, we permeabilized lysosomal membranes using the lysosomotropic agent Leu-Leu-O-Me (LLOMe) and confirmed membrane damage with the marker galectin-3^(51)^ (Fig. S4C). Elevated lysosomal cholesterol was detected in the LLOMe treatment as well as in the treatment by lysosome deacidifying reagents, such as the ionophore monensin or V-ATPase inhibitors bafilomycin and chloroquine (Fig. S4D). These results showed that lysosome damage and pH increase could sequestrate cholesterol. However, we did not detect galectin-3 puncta in the Flp-in cells (Fig. S4E), suggesting that ORF3a does not markedly damage lysosomes in these cells, and that there are other mechanisms that interrupt cholesterol independent of lysosome damage.

We next examined the lysosomal cholesterol transporters NPC proteins in the Flp-In cells. Immunoblotting the endogenous proteins did not show an obvious difference in NPC protein levels between the four Flp-in cell lines (Fig. S4F) or between mock and infected cells (Fig. S4G). Immunofluorescence confocal microscopy revealed that NPC1 colocalized with LAMP2 (Fig. 4E), and no difference of Pearson correlation coefficient was detected by high-content imaging quantification (Fig. 4F). In contrast, a mislocalization of NPC2 from the lysosomes was observed in the CoV-2-ORF3a cells (Fig. 4G), reproduced in a quantitative measurement of Pearson correlation coefficient between NPC2 and LAMP1 (Fig. 4H). This mislocalization was largely rescued by the W193A mutant (Fig. 4G,H). Further analysis of protein secretion revealed an increased level of NPC2 and another lysosomal lumenal protein, cathepsin D, in the culture media of CoV-2-ORF3a cells (Fig. 4I,J). These results motivated us to test whether lysosomal transmembrane proteins, such as LAMPs, can localize to the lysosomes. Distinct from lysosomal transmembrane proteins that are transported from TGN to lysosomes through biosynthesis pathway, Ragulator complex is associated with lysosome membrane through lipidation of its subunit LAMTOR1, which scaffolds other subunits. By measuring the colocalization between LAMP2 and the Ragulator subunit LAMTOR4, we did not see a difference between the Flp-In cells (Fig. S4H,I). Together, these results indicated that ORF3a induces a trafficking defect of lysosome lumenal proteins, likely due to a dysregulation of protein transport between TGN and endosomes.

### ORF3a disturbs the endosome-to-TGN transport

The NPC2 trafficking defect observed in CoV-2-ORF3a cells resembled what was observed in VPS39-deleted cells in our previous study, in which the sorting receptor for lysosomal lumenal proteins, CI-MPR, is abnormally degraded in lysosomes^(18)^. In CoV-2-ORF3a cells, CI-MPR protein level did not reduce (Fig. S4F), but it dispersed from its usual location near the microtubule-organizing center (MTOC) to the cell periphery (Fig. 5A). In addition, a more severe dispersion of the TGN protein TGN46 was observed in CoV-2-ORF3a cells (Fig. 5A). The positioning changes of CI-MPR and TGN46 was confirmed by high-content imaging analysis (Fig. 5B,C), in which cellular area between MTOC and cell boundary was divided into “center”, “middle”, and “periphery” to reflect protein cellular distribution^(52)^. We also examined Golgi matrix protein 130 (GM130) but did not see a dispersion (Fig. S5A), indicating that the dispersion of CI-MPR and TGN46 in the Flp-in cells was not due to Golgi fragmentation^(53,54)^}. The dispersed CI-MPR and TGN46 were partially colocalized with LAMP2 (Fig. 5A, arrows), and quantitative results showed an increased Pearson correlation coefficient between CI-MPR and LAMP2 (Fig. 5D). Consistently, CoV-2-ORF3a cells had an increased localization between TGN46 and Rab7 (Fig. 5SB,C). These results indicate that CI-MPR and TGN46 were retained within the Rab7-or LAMP2-positive endosomes and lysosomes in the CoV-2-ORF3a cells, suggesting that TGN-to-endosome transport was increased, and/or retrieval from endosomes and lysosomes was decreased, resulting in protein prolonged retention within the endosome-lysosome system.

**Figure 5.**
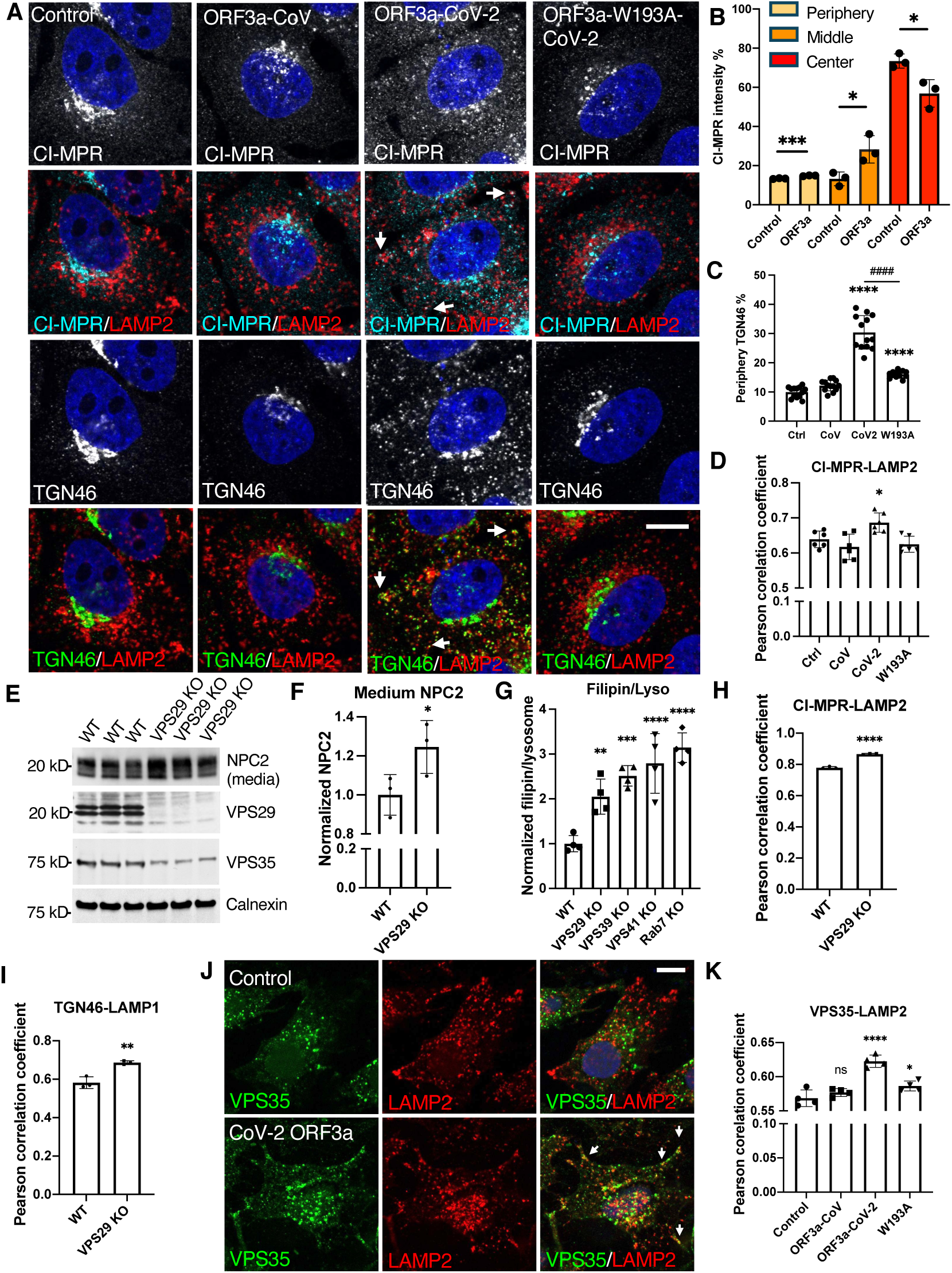
Decreased endosome-to-TGN trafficking. **A-D**, HeLa Flp-In cells were fixed after 16-h induction of protein expression and immunostained with the indicated antibodies. Confocal microscopy (images) and high-content imaging (bar charts) were performed to analyze protein cellular distribution and colocalization. Periphery was defined as a ring area shrunk from cell boundary with a gap distance 15 pixels (B,C). Center was defined as a dilated circle from nuclear boundary with a distance of 20 pixels (B). Middle was defined as the area between periphery and center (B). **E,F**. Wildtype and VPS29-KO HeLa cell were cultured in serum-free media for 6 hours, and the media were collected before cells were lysed. The media and cell lysates were subjected to immunoblotting with the indicated antibodies. **G**. Vesicular filipin was measured by high-content imaging and normalized to lysosomes in the indicated cells. **H-K**, Cells were fixed and immunostained with the indicated antibodies. Confocal microscopy (images) and high-content imaging (bar charts) was performed to analyze protein colocalization. Bar graphs are presented as mean ± SD. *p* values were determined using *t’* test or One-way ANOVA test *, *p*<0.05, **, *p*<0.01, \*\*\**, p*<0.001, **** (vs. control) or #### (C, CoV-2 vs W193A), *p*<0.0001, n.s., not significant. Scale bars, 5 μm. Control (Ctrl), linker peptide. CoV, SARS-CoV ORF3a. CoV-2, SARS-CoV-2 ORF3a. W193A, SARS-CoV-2 ORF3a-W193A mutant.

Retrieval of endosomal proteins relies on the retromer^(16,55)^ and retriever^(56)^ complexes. KO of their shared subunit VPS29 resembled the defects observed in CoV-2 ORF3a cells, including increased NPC2 secretion (Fig. 5E,F), lysosomal cholesterol accumulation (Fig. 5G), and enhanced colocalization of CI-MPR and TGN46 with LAMP1 (Fig. 5H,I). These results indicate that disruption of endosomal protein retrieval impairs lysosome cholesterol egress. We next examined the Flp-In cells and found that the retromer subunit VPS35, which mediates endosome-to-Golgi transport, exhibited increased colocalization with LAMP2 in CoV-2-ORF3a cells (Fig. 5J, arrows). High-content imaging quantification showed an elevated Pearson’s correlation coefficient between VPS35 and LAMP2, which was rescued in W193A cells (Fig. 5K). Together, these findings indicate that the ORF3a-VPS39 interaction disrupts endosome-to-TGN transport by retaining retromer in LAMP2-positive endosomes/lysosomes, resulting in reduced recycling of CI-MPR and targeting of NPC2 to lysosomes.

### ORF3a reduced BMPs

BMPs, localized exclusively on late endosome and lysosome membranes, physically interacts with NPC2 to facilitate cholesterol egress^(33,34)^. Using an antibody against BMPs^(26)^, we observed a significant reduction in BMP levels in SARS-CoV-2-infected Vero E6 cells, starting 12 hours post-infection (Fig. 6A,B). To test if this alteration was related to ORF3a, we examined the Flp-In cells and found that the BMP level was reduced by 20% in CoV-2-ORF3a cells, compared to the control cells. This reduction was partially rescued in W193A cells, while the CoV-ORF3a cells exhibited no change (Fig. 6C,D). Shotgun lipidomics of total lipids extracted from the Flp-In cells confirmed these results: the two most abundant BMP species, 18:1-18:1 and 18:1-22:6, were reduced by approximately 20% in CoV-2-ORF3a cells and restored in W193A cells (Fig. 6E). These results indicate that CoV-2-ORF3a alone can decrease BMP levels, which is dependent on the ORF3a-VPS39 interaction, aligned with the observed cholesterol sequestration. Furthermore, exogenous BMP treatment for 2 hours in serum-free media decreased the lysosomal cholesterol levels by 25% in CoV-2-ORF3a cells (Fig. 5F). Together, these results suggest that reduced BMP is an additional mechanism underlying SARS-CoV-2-induced cholesterol sequestration.

**Figure 6.**
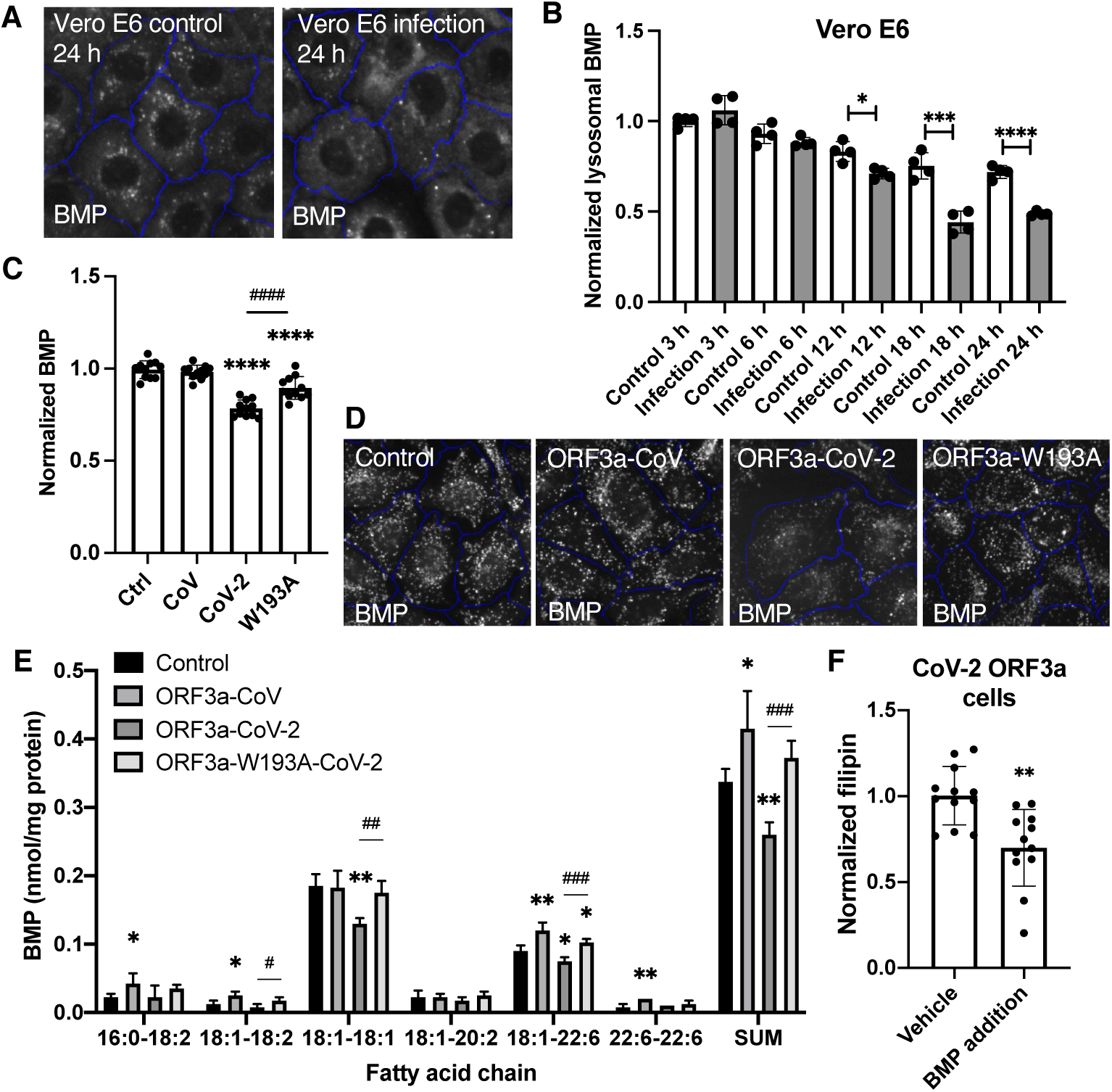
Analysis of BMP levels. **A,B.** Vero E6 cells were infected with SARS-CoV-2, fixed at the indicated time points post-infection, and immunostained with BMP and LAMP1 antibodies. LAMP1-positive BMP puncta were quantified with the high-content imaging system. Representative images were shown in B. **C,D.** BMP in the HeLa Flp-In cells were measured as described in A. **E.** The indicated cells were harvested and lysed for total lipid extraction. Shotgun lipidomics was performed to identify and quantify bio(monoacylglycero)phosphate (BMP) species based on the fatty acid chains, labeled as carbon number: double-bond number. BMP was normalized to protein concentrations. **F.** Cells were incubated with serum-free media containing 1% BSA and 1 µM BMP for 2 hours, fixed and stained with filipin. filipin levels were measured with high-content imaging. Bar graphs are presented as mean ± SD. *p* values were determined using *t’* test or One-way ANOVA test *, *p*<0.05. **, ##, *p*<0.01. \*\*\**, p*<0.001. ****, ####, *p*<0.0001, n.s., not significant. Scale bars, 5 μm. Control (Ctrl), linker peptide. CoV, SARS-CoV ORF3a. CoV-2, SARS-CoV-2 ORF3a. W193A, SARS-CoV-2 ORF3a-W193A mutant.

### ORF3a interrupted lysosome-mitochondrion interaction

To investigate the mechanism underlying BMP reduction, we examined key enzymes involved in BMP biosynthesis and turnover, including CLN5^(37)^, PLD3, PLD4^(38)^, ABHD6^(35)^, and PLA2G15^(39)^. With the exception of PLA2G15, which did not display a punctate pattern upon transfection, we did not observe a reduced colocalization with lysosomes (Fig. S6A-H) or reduced expression levels (Fig. S6I) of these proteins, suggesting these enzymes are not responsible for the BMP reduction in CoV-2-ORF3a cells.

We then isolated the lysosomes by Lyso-IP^(57)^ followed by mass spectrometry of lysosomal proteomes from the control, CoV-2-ORF3a and W193A Flp-In cells (Tab. 1). Comparison revealed 277 proteins uniquely present in CoV-2-ORF3a samples (gained, Tab. 2), and 348 proteins increased at least 2-fold relative to W193A cells (increased, Tab. 3). In contrast, 160 proteins were undetected (lost, Tab. 2), and 87 proteins decreased by ≥50% in CoV-2-ORF3a samples (decreased, Tab. 3). Gene ontology analysis using DAVID Bioinformatics indicated that the gained and increased proteins largely originated from cytoplasm (32%) and nuclei (25%) (Fig. 7A), while the lost and decreased proteins were predominantly mitochondrial (45%) and ER-derived (21%) (Fig. 7B,C,D). Among the top 20% of lost proteins, 43.7% proteins were mitochondrial, including outer membrane, inner membrane, and matrix proteins (Fig. 7C). Similarly, in the reduced proteins, the majority was from mitochondria, regardless their mitochondrial localization (Fig. 7D). These results suggest a reduced lysosome-mitochondrion interaction in CoV-2-ORF3a cells.

**Figure 7.**
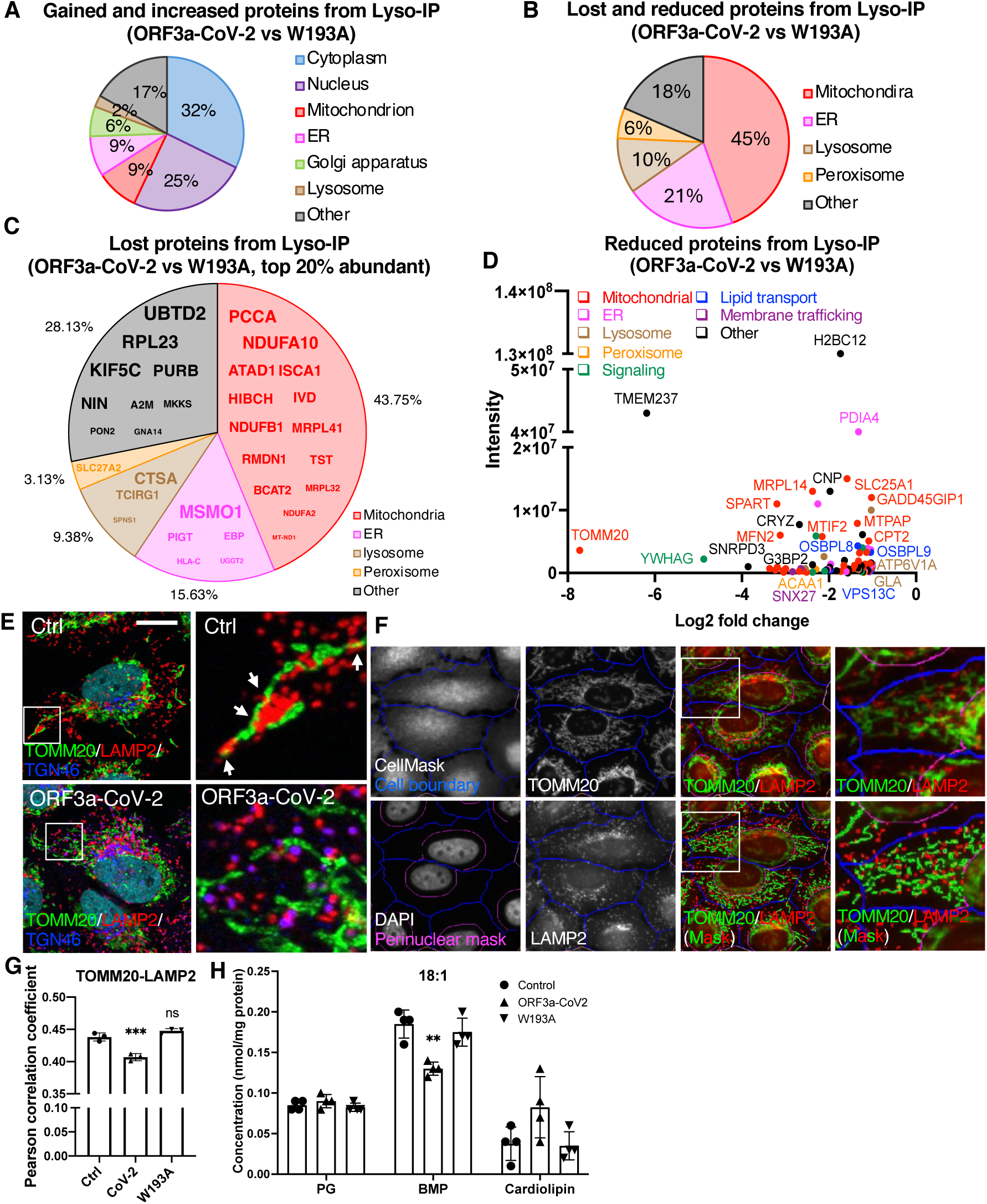
Decreased lysosome-mitochondrion interaction. **A-D.** Lysosomes were isolated from ORF3a and W193A Flp-In cells, and the total protein were extracted for mass spectrometry analysis. The proteins only presenting in ORF3a lysosomes (gain) or increased more than 2-folds compared to W193A lysosomes (increased) are shown in the pie chart based on their cellular localization (**A**). The proteins only presenting in W193A lysosomes (lost) or decreased more than 50% compared to W193A (reduced) are shown in **B.** Amount the lost proteins, as defined above, top 20% abundant proteins in W193A lysosomes are shown in **C.** The reduced proteins, as defined above, were plotted in **D**. The color codes reflect their cellular localization or functions. **E**. The control and CoV-2 ORF3a cells were fixed and immunostained with the indicated antibodies. Confocal microscopy was applied for imaging. Arrows indicate the lysosome-mitochondrion physical interactions. Scale bar, 5 μm. **F,G.** The Flp-In cells were fixed and immunostained with the indicated antibodies accompanied with CellMask and DAPI. High-content imaging system was used for image collection and analysis. Cell boundary and perinuclear area were defined based on CellMask and DAPI, respectively. Lysosome and mitochondrial masks were defined with LAMP2 and TOMM20 signals, respectively. The colocalization between lysosomes and mitochondria were quantified and presented as Pearson correlation coefficient. **H.** Total lipids were extracted from the indicated cells and subjected to shotgun lipidomics analysis. PG, BMP, and Cardiolipin with the 18:1 fatty acid chains are presented. PG, phosphatidylglycerol. BMP, bis(monoacylglycerol)phosphate. Bar graphs are presented as mean ± SD. *p* values were determined using One-way ANOVA test **, *p*<0.01. \*\*\**, p*<0.001. n.s., not significant.

Confocal imaging showed frequent lysosome–mitochondria interactions in control cells (Fig. 7E, upper, arrows), while CoV-2 ORF3a cells exhibited fewer physical contacts, though organelles remained proximal (Fig. 7E, lower). To quantify the physical interactions, we defined the cytoplasm area in each cell and excluded the perinuclear clouds to reduce the false overlap due to a usual crowdedness of organelles at that area (Fig. 7F). Indicated by Pearson correlation coefficient, the interactions between mitochondria and lysosomes was significantly decreased in the CoV-2-ORF3a cells and was rescued in the W193A cells (Fig. 7G). High-content imaging of TOMM20 revealed slightly increased mitochondrial abundance and similar morphology in both CoV-2-ORF3a and W193A cells (Fig. S7A,B,C), confirming that the reduced lysosome-mitochondrion interaction was not due to fewer mitochondria or altered mitochondrial morphology.

We hypothesized that impaired mitochondrion-lysosome interaction may limit mitochondrial transportation of PGs, the precursors for BMP biosynthesis. Lipidomics revealed unchanged PG levels among the three cell types, but mitochondrial cardiolipin, a PG derivative, increased in CoV-2 ORF3a cells (Fig. 7H), suggesting that interrupted PG exportation from mitochondria resulted in an increased local PG conversion and decreased BMP synthesis in lysosomes.

### ORF3a disturbed mitochondrion-lysosome membrane contact sites

The direct physical interactions between mitochondria and lysosomes can be achieved through autophagy, mitochondrion-derived vesicles (MDVs), and mitochondrion-lysosome membrane contact sites (MCS). It was shown that autophagy flux is interrupted by ORF3a-VPS39 interaction, which inhibits HOPS-mediated fusion between autophagosomes and lysosomes^(32)^. We observed the same defect shown as increased LC3-II levels in CoV-2-ORF3a cells (Fig. S8A). However, inhibiting autophagy fusion by STX17 KO (Fig. 8A), or interrupting earlier autophagy steps by knockdown (KD) of ATG5, ATG7, or ULK1 (Fig. 8C) did not alter BMP levels (Fig. 8B,D). These results excluded autophagy as a PG transport pathway.

**Figure 8.**
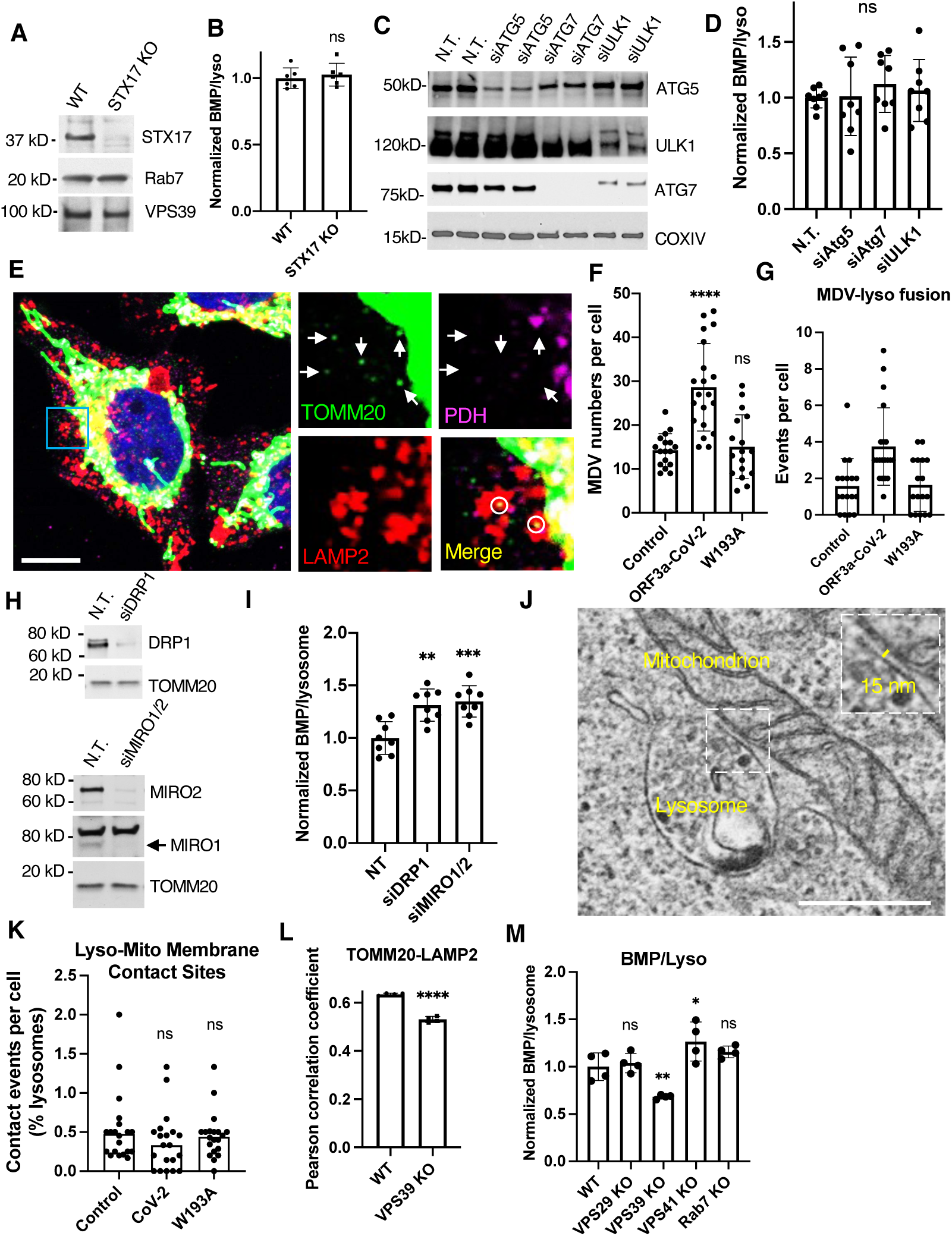
Involvement of impaired lysosome-mitochondrion membrane contact sites in BMP reduction. **A.B**, Wildtype and STX17-knockout (KO) HeLa cells were lysed for immnoblotting with the indicated antibodies (**A**) or fixed for immunostaining and hign-content imaging quantification (**B**). **C.D**, Wildtype HeLa cells were transfected with siRNA targeting the indicated autophagy genes. Non-targeting siRNA (N.T.) served as a control. Protein levels were examined by immunoblotting (**C**), and the BMP levels were measured by immunostaining followed by high-content imaging (**D**) at 2 days post-transfection. **E-G**, The Flp-In cells were fixed and immunostained with a TOMM20 (marker of mitochondrion-derived vesicles, MDVs), PDH (maker of mitochondria but not MDVs), and LAMP2 (lysosome marker) antibodies. Confocal microscopy was performed to identify MDVs (arrows) and the MDVs fused with lysosomes (circles) (**E**). The MDV numbers (**F**) and the number of lysosome-fused MDVs (**G**) were quantified using FIJI. Scale bar, 5 μm. **H.I**, HeLa cells were transfected with the siRNAs of N.T., DRP1, or combined MIRO1 and MIRO2 and subjected to immunoblotting (**H**) or high-content imaging quantification (**I**). **J.K**, The Flp-In cells were seeded on glass coverslips, fixed, and subjected to electron microscopy. Randomized cells were imaged at mitochondrion-existing areas. Lysosome-mitochondrion membrane contact sites were identified by their membrane structures and the distance between membranes (**J**) and quantified from 20 cells of each type (**K**). Scale bar, 0.5 μm. **L.M**, Wildtype and VPS39-knockout (KO) HeLa cells were immunostained and subjected to high-content imaging quantification. Bar graphs are presented as mean ± SD. *p* values were determined using *t’* test or One-way ANOVA test. *, *p*<0.05. **, *p*<0.01. \*\*\**, p*<0.001. ****, *p*<0.0001. n.s., not significant.

We next tested MDVs, identified by a combined staining of the MDV marker TOMM20 and a mitochondrial but not MDV protein, pyruvate dehydrogenase (PDH) (Fig. 8E, arrows). We also included a lysosome marker to visualize the fusion between MDV and lysosomes (Fig. 8E, circles). CoV-2-ORF3a cells exhibited more MDVs and MDV-lysosome fusion events (Fig. 8F,G), which did not align with the BMP reduction (Fig. 6C). Furthermore, KD of MDV biogenesis genes, DRP1 or MIRO1/2^(58)^ (Fig. 8H), did not decrease BMP levels (Fig. 8I), indicating that MDVs do not mediate PG delivery to lysosomes.

Last, we applied electron microscopy to assess mitochondrion-lysosome MCS. We randomly chose cells at low magnification and image areas containing abundant mitochondria at higher magnification. Lysosome-to-mitochondrion ratios were similar across cell types (Fig. S8B), but MCS events, identified by intact membrane within 10 to 30 nm distance (Fig. 8J), decreased in CoV-2-ORF3a cells and were rescued in the W193A cells, though it was not statistically significant (Fig. 8K). VPS39 is known to mediate the MCS formation in a HOPS-independent manner in yeast^(59)^. Consistently, VPS39 KO decreased TOMM20-LAMP2 overlap and BMP level, whereas VPS29, VPS41, or Rab7 KO did not (Fig. 8L,M). There findings suggest that ORF3a inhibited VPS39-mediated lysosome-mitochondrion MCS formation and impaired PG transportation, consequently reducing BMP synthesis.

## DISCUSSION

Our study identifies SARS-CoV-2 ORF3a as a potent disruptor of lysosomal cholesterol transport through dual interference with NPC2 trafficking and BMP biogenesis. By binding VPS39, ORF3a blocks retromer-mediated retrieval of CI-MPR and reduces mitochondrion-to-lysosome lipid transfer, thereby sequestering cholesterol within lysosomes. These findings expand our understanding of cellular cholesterol regulation and the functions of HOPS and VPS39, point to a regulatory pathway for BMP biosynthesis, and reveal a viral strategy that links lysosomal lipid metabolism to SARS-CoV-2 pathogenesis.

Cholesterol is not only a structural component of cellular membranes but also a dynamic signaling molecule whose intracellular distribution dictates multiple metabolic pathways. Our results demonstrate that SARS-CoV-2 impedes lysosomal cholesterol egress (Fig. 1), adding to the growing recognition that compartmentalized cholesterol pools, rather than bulk cholesterol, drive cellular physiology^(60,61)^. ORF3a’s ability to retain cholesterol within lysosomes (Figs. 2&4C,D) reveals a new viral strategy to manipulate host lipid homeostasis and may explain systemic lipid alterations in COVID-19 patients^(1–5)^. More broadly, our findings place lysosomal cholesterol transport at the crossroads of host–pathogen interactions and suggest that this pathway may represent a therapeutic vulnerability.

BMPs play essential roles in lysosomal biology and have been implicated in a wide range of human diseases^(27,28)^, such as infectious disease^(30,32,62,63)^, and neurodegenerative disorders^(64,65)^. Here, we demonstrate that SARS-CoV-2 and its ORF3a protein reduce BMP levels (Fig. 6A-E), and that restoring BMP partially rescues cholesterol egress (Fig. 6F). This finding highlights BMP as a limiting factor for lysosomal cholesterol export, in agreement with prior studies^(33,33,34)^. Notably, ORF3a alters PG metabolism by decreasing its lysosomal products (BMPs) and increasing its mitochondrial products (cardiolipins) (Fig. 7H). This shift uncovers a previously unrecognized regulatory mechanism in BMP homeostasis, the supply of mitochondrial precursors. While BMP levels were thought to be controlled primarily by lysosomal synthetase and hydrolases^(35,37–39)^, our data suggest that cross-organelle lipid transfer provides an additional layer of regulation. Importantly, by testing multiple possible routes, we found that mitochondrion-lysosome membrane contact sites likely serve as the pathway for PG transport, rather that autophagy or mitochondrial-derived vesicles (Fig. 8). These finding establish BMP regulation as a point of crosstalk between mitochondrial phospholipid metabolism and lysosomal cholesterol transport, with broad implications for both cellular physiology and viral pathogenesis.

The canonical view of HOPS has centered on its role as a tethering complex that mediates late endosome–lysosome and autophagosome–lysosome fusion^(17)^. By leveraging the VPS39–ORF3a interaction and the W193A mutant, our study expands this paradigm by showing that VPS39 also regulates two distinct aspects of lysosomal activities, regulation of retromer-dependent trafficking^(66)^ (Fig. 5) and formation of mitochondrion–lysosome membrane contact sites necessary for BMP synthesis (Fig. 8J-M). These findings position VPS39 as a multifunctional coordinator at the lysosome, integrating protein trafficking and lipid exchange. The dual disruption of these VPS39-dependent functions by ORF3a reframes HOPS/VPS39 not solely as a fusion mediator, but as a broader regulatory hub essential for lysosomal homeostasis.

Our study establishes ORF3a as a viral effector that directly manipulates host lipid metabolism. Unlike SARS-CoV ORF3a^(67)^, its SARS-CoV-2 counterpart acquired the ability to bind VPS39^(43,44,68)^, suggesting an evolutionary adaptation to exploit lysosomal cholesterol regulation. Since plasma membrane cholesterol promotes SARS-CoV-2 entry^(11,12)^, restricting cholesterol export from lysosomes could reduce cholesterol at the plasma membrane and thereby limit the viral entry. Consistent with this model, pharmacological blockade of NPC1 with U18666A sequesters cholesterol in lysosomes and decreases SARS-CoV-2 infection (Fig. S8C–E). In addition, BMPs are required for viral fusion within endosomes/lysosomes^(63)^, and reduced BMP levels may impair this step of the viral life cycle. Importantly, ORF3a is an accessory protein that is not incorporated into virions but translated only after infection from viral subgenomic RNAs^(69)^. Thus, its interference with cholesterol export and BMP synthesis is unlikely to affect the current infection but may restrict secondary infection events. Furthermore, ORF3a’s previously reported activities, including increasing lysosomal pH and damaging lysosomal membranes, may promote lysosome exocytosis, a route for viral egress^(42,70)^. Taken together, these findings suggest that SARS-CoV-2 use ORF3a to modulate multiple lysosomal functions, integrating cholesterol trafficking, lipid metabolism, and membrane dynamics to optimize different stages of its life cycle.

Our study has several limitations. First, although we demonstrate ORF3a-induced cholesterol sequestration and identify VPS39-dependent mechanisms, the downstream consequences for viral replication, immune responses, and cell survival remain to be fully elucidated *in vivo*. Second, while we observed reduced mitochondrion–lysosome contacts and altered BMP levels, the molecular intermediates mediating phosphatidylglycerol transfer remain undefined. Finally, the interplay between cholesterol sequestration and systemic lipid dysregulation observed in COVID-19 patients requires further integration with clinical data. Notably, SARS-CoV-2 infects hepatocytes^(71)^, and given the conserved role of HOPS in lysosomal trafficking, similar lipid defects may occur in the liver, a central regulator of cholesterol and lipoprotein metabolism. Future work will therefore examine hepatocytes to determine whether ORF3a-mediated lysosomal disruptions contribute to the lipoprotein abnormalities observed in COVID-19 and long COVID.

In summary, this work uncovers cholesterol transport and BMP biogenesis as dual regulatory nodes governed by VPS39 and reveals how SARS-CoV-2 ORF3a exploits these pathways to rewire lysosomal lipid metabolism. These findings advance fundamental understanding of lysosomal protein trafficking and lysosome–mitochondrion crosstalk, broaden the functional repertoire of HOPS/VPS39, and provide new perspectives on viral pathogenesis and cholesterol-related diseases.

## MATERIALS AND METHODS

### Cells and virus

A549 and HeLa cells were cultured in Dulbecco’s Modified Eagle’s Medium (DMEM) supplemented with 10% heat-inactivated fetal bovine serum (FBS), 25 mM HEPES, and MycoZap Plus-CL (Lonza, Basel, Switzerland) at 37°C with 5% CO_2_. Vero E6 cells (ATCC, CRL-1586) were cultured in DMEM supplemented with 10% heat-inactivated FBS, 1% pen/strep, 2mM L-glut, 1% non-essential amino acids, 1% HEPES. Severe acute respiratory syndrome coronavirus 2 (SARS-CoV-2, Isolate USA-WA1/2020) was acquired from BEI Resources (NR-52281) and propagated in Vero E6 cells for 3 days. Infectious virus was isolated by harvesting cellular supernatant and spinning at 1000 g for 10 minutes to remove cellular debris and stored at –80°C. SARS-CoV-2 infectious particles were quantified in media by standard plaque assay^(72)^

### SARS-CoV-2 infections

Coverslips or optic 96-well plates (Agilent, Santa Clara, CA) were coated with collagen solution (0.03-0.04 mg/ml collagen in 0.02 N acetic acid) for > 30 min and then washed twice with PBS before receiving cells. A549-hACE2 cells were seeded on coverslips at 0.1 × 10^6^ cells per well or in optical 96-well plates at 0.012 × 10^6^ cells per well and infected with SARS-CoV-2 in DMEM (4% FBS) at MOI: 4 and incubated at 37°C for 1 hr on the next day. Cells were then washed twice with 1x PBS and normal culture media was added. At the indicated times post-infection, cells were fixed with 4% PFA. Vero E6 cells were seeded in optical 96-well plate around 0.025 × 10^6^ cells per well and infected with SARS-CoV-2 at MOI: 0.1 in 2% FBS media. The control group of cells were incubated with the media only.

### Plasmid and siRNA transfection

Cells were seeded a day before transfection and receive plasmids mixed with transfection reagent Lipofectamine 2000 (Thermo Fisher Scientific) or TransIT-LT1 (Mirus Bio, Madison, WI), following the manufactures’ instructions. Cells were analyzed at 24-or 48-h post transfection. On-TARGETplus SMARTpool siRNAs were applied (Horizon Discovery, Cambridge, UK) for two-shot transfection at Day 1 and Day 3 after seeding. The transfection reagents Lipofectamine 2000 (Thermo Fisher Scientific) or TransIT-X2 (Mirus Bio) were used to deliver siRNA, and cells were analyzed 6-7 days after seeding.

### CRISPR KO

Genes were edited using the CRISPR/Cas9 system. Guide RNAs (gRNAs) were designed using Benchling () Four to six 20-base pair (bp) gRNAs were selected and tested for the cleavage in pilot experiments. gRNA was introduced into the px458 plasmid containing GFP sequence (Addgene, Cambridge, MA). Cells were transfected with the plasmids, and GFP-positive cells were collected by cell sorter 48 h after transfection. Transformants were kept in normal medium for another 12 days to allow single colony formation. Genomic DNA was extracted from individual colonies, and cleavage of the target sequence was tested by PCR followed by Sanger sequencing and immunoblotting. Three to six single colonies were pooled for experiment use.

Guide RNA target sequences used in this study:

Rab7 KO: TAGTTTGAAGGATGACCTCT and TTGCTGAAGGTTATCATCCT

VPS29 KO: GCAAACTGTTGCACCGGTGT and CATAACTCTCTTTGGTGCAA

Stx17 KO: TGGAGAAGACAGCTGTTACCAGGG and TTAAGATAGTAATCCCAACAGACC

### Generation of flip-in cells

HeLa Flp-In host cells (Thermo Fisher Scientific, Waltham, MA) were transfected with the plasmids encoding FLAG-tagged proteins or linker peptide and subjected to hygromycin selection after 48 h post transfection in DMEM as described above but replacing regular FBS with tetracycline-free FBS (Thermo Fisher Scientific) (tet-free media). Following 2-week selection, transformants were kept in tet-free medium for another 12 days to allow single colony formation. Protein expression was induced by addition of 1 µg/ml doxycycline for 16 hours. The colonies had relatively low expression levels of FLAG were chosen and pooled for experiments, determined by immunoblotting of FLAG.

### Immunofluorescence, Filipin staining, and confocal microscopy

Fixed cells were washed 3 times with PBS, and primary antibodies were diluted in 1% BSA in PBS supplemented with 0.2% saponin and applied to cells for 1 h at 37°C or overnight at 4°C. Alexa-conjugated secondary antibodies (Thermo Fisher Scientific) were applied with the same saponin-contained BSA-PBS solution for 30 min at 37°C. Coverslips were mounted with DAPI-contained mounting reagents (Electron Microscopy Sciences, Hatfield, PA) and allowed to dry at 37°C for at least 1 h. To stain free cholesterol, 25 µg/ml filipin complex (Millipore Sigma, St. Louis, MO) in PBS was applied to fixed cells at 37°C for 30 min. When immunostaining was performed with filipin, primary antibodies and Alexa-conjugated secondary antibodies along with DRAQ7 (Novus Biologicals, Centennial, CO) were sequentially applied to filipin-stained cells without detergent permeabilization. Then cells were mounted with mounting reagents without DAPI (Electron Microscopy Sciences) at 37°C for at least 1 h. Images were acquired using a Zeiss LSM800 confocal microscope equipped with the software ZEN (Zeiss, Oberkochen, Germany).

### High-content imaging and image analysis

Cells in optical 96-well plates were fixed and stained as described above and maintained in PBS for immediate imaging. Imaging and analysis were performed using the CellInsight Microscope platform (Thermo Fisher Scientific). DAPI-or DRAQ7-stained nuclei were scanned to determine focus through the programmed autofocus function. Cytoplasmic signals from HCS CellMask Near-IR Stain (Thermo Fisher Scientific) were applied to define individual cell boundaries. Cells were excluded based on criteria including incomplete cell boundaries at image edges, abnormal size or shape, and signs of cell death.

For cellular punctum analysis, the cell boundary-enclosed areas are defined as ROI_A. Filipin, LAMP, or BMP puncta were detected as spots (2 pixels in size) using the software “spot detection” function. Line-shaped filipin signals from the plasma membrane were excluded by setting the validation criteria for shape. Puncta that overlapped with ROI_A were quantified by their fluorescence intensity. To measure colocalized signals, the software function of “colocalization” was applied, and the output feature “ROI_A(B) Correlation Coefficient” was extracted for the quantification of Pearson Correlation Coefficient.

To analyze the infected cells, fixed cells were immunostained with an dsRNA antibody, and its signals were used to set the threshold for detection of infected cells, using the function “Reference” of the software.

### Sample Processing for electron microscopy

Cells on glass coverslips were fixed in 0.1 M PIPES buffer containing 3% formaldehyde, 2% glutaraldehyde, and 1.5 mM CaCl2, then washed in several buffer changes before secondary fixation in 0.1 M PIPES containing: 1% osmium tetroxide for 15 min, 5 min PIPES rinse, 15 min 1% carbohydrazide in PIPES, 5 min PIPES rinse, and 15 min 1% osmium tetroxide (OCO). Samples were then washed with buffer, followed by ultrapure water, before being dehydrated through a graded series of ethanol to propylene oxide, infiltrated with Epon-Araldite resin, embedded, and heat-cured. The glass was removed from the cured resin blocks so the cells could be re-mounted for en face sectioning. Thin sections (60-80 nm) were mounted on Cu grids, post-stained with uranyl acetate and Reynold’s lead citrate, and examined in a Hitachi HT7700 TEM operated at 80 kV. Images were captured with an AMT XR-81 CCD camera.

### Cholesterol measurement by GC-MS

Cells were collected in PBS with a cell scraper after washing twice with PBS, and the total lipids were extracted by adding lipid-extraction solvent (chloroform:methanol:acetic acid = 50:50:1) of 90% volume of PBS. The organic phase and aqueous phase were separated by 10 minute-centrifugation at 21,000 g. Organic phase was collected and transferred into a new tube and dried with vacuum. The extracted lipids were stored at –80°C for further analysis. Cholesterol calibration standard solutions were prepared from the concentration of 50 µg/ml to 400 µg/ml along with 80 µg/ml 5a-cholestane as the internal standard. The linear calibration curve was obtained with R2 =0.9884. The dried lipids were dissolved with hexane and mixed with 4 µg 5a-cholestane. The GC-MS analysis was conducted on Agilent 6890/5973 with selected ion monitoring (SIM) mode.

### Shotgun Lipidomics

Lipid species were analyzed using a multidimensional mass spectrometry-based shotgun lipidomics approach^(73)^. In brief, each cell sample homogenate containing 0.3 mg of protein which was determined with a Pierce BCA assay was accurately transferred to a disposable glass culture test tube. A premixture of lipid internal standards was added prior to conducting lipid extraction for quantification of the targeted lipid classes. Lipid extraction was performed using a modified Bligh and Dyer procedure (Wang and Han 2014), and each lipid extract was reconstituted in chloroform:methanol (1:1, v:v) at a volume of 400 µL/mg protein. Derivatization of lipid extracts for analysis of free fatty acid (FFA) and bis(monoacylglycero)phosphate (BMP) was performed as described previously^(74,75)^ before lipidomics analysis. Lyso-phosphatidylglycerol (LPG) in the aqueous phase was enriched using a HybridSPE cartridge. After washing with methanol, lysophospholipids were eluted with methanol/ammonia hydroxide (9:1), dried and reconstituted in methanol for lipidomics analysis ^(76)^. For shotgun lipidomics, individual lipid samples prepared as aforementioned was further diluted to a final concentration of ∼500 fmol total lipids per µL. Mass spectrometric analysis was performed on a triple quadrupole mass spectrometer (TSQ Altis, Thermo Fisher Scientific, San Jose, CA) and a Q Exactive mass spectrometer (Thermo Scientific, San Jose, CA), both of which were equipped with an automated nanospray device (TriVersa NanoMate, Advion Bioscience Ltd., Ithaca, NY) as described^(77)^. Identification and quantification of lipid species were performed using an automated software program^(78,79)^. Data processing (e.g., ion peak selection, baseline correction, data transfer, peak intensity comparison, and quantitation) was performed as described^(79)^. The results were normalized to the protein content (nmol lipid/mg protein).

### Analysis of intracellular and extracellular proteins by Western blotting

To detect secreted NPC2 and cathepsin D, cells were kept in a minimum volume of DMEM without serum for 6 h after three-time washing with the same medium. The media were collected and centrifuged at 1,000 g for 5 min to pellet any detached cells. The supernatants were mixed with SDS loading buffer and subjected to SDS-PAGE and immunoblotting. After collecting the media, cells were incubated in complete DMEM for 2 h. Cell lysates were obtained by using 2X SDS sample buffer.

### Quantification and statistical analysis

Image J was used for quantifying fluorescence signals of randomly-taken images from at least three independent experiments. The signal intensity of vesicular filipin was measured using the function of “Analyze particles”, and co-localization was analyzed with the plug-in “PSC Colocalization”^83^. High-content imaging results were obtained from more than 1,000 cells per well and at least 3 wells per experiment. The number of trials is specified in the legends where appropriate. The bar graphs show the pooled results displayed as mean ± SD. Colocalization results include frequency distribution. Statistical significance was determined by comparing two datasets using a *t* test with one-tailed distribution or more than two datasets using one-way ANOVA, using the software Prism 9 (GraphPad). Probability values and number of trials are given in the figure captions and the legends where appropriate. The statistical significance is generally denoted as follows: *, *p* < 0.05, **, *p* < 0.01, ***, *p* < 0.001, ****, *p* < 0.0001, and n.s., not significant.

## AUTHOR CONTRIBUTIONS

J.P. conceived the project. B.G. and J.P. performed most of the experiments. V.M.V. characterized VPS29-KO cells and performed secretion assays. O.K. quantified NPC2-LAMP1 colocalization. R.L. measured total cholesterol levels by GC-MS. J.J. made SARS-CoV-2 ORF3a cell line. M.I. made VPS29-KO cells. H.W. and X.H. performed lipidomics analysis. A.K. performed SARS-CoV-2 infection in A549 cells. C.Y. and S.B. performed the infection in Vero E6 cells. M.R.L. analyzed the structures of SARS-CoV and SARS-CoV-2 ORF3a. B.G. and J.P. analyzed the data. J.P. wrote the manuscript. All the authors edited the manuscript.

## ACKNOWLEDGMENTS

We thank A. Kajon for gift of A549-hACE2 cells, P. Lobel for mch-NPC2 construct and NPC2 antibody, J. Bonifacino for VPS29-KO HeLa cells, and J. Anderson for experimental assistance. EM data were generated in the HSC-Electron Microscopy Facility, supported by University of New Mexico Health Sciences Center (HSC). This research made use of the CellInsight Microscope Platform and confocal microscope from Autophagy, Inflammation, and Metabolism Center of Biomedical Research Excellence, funded by NIGMS, NIH (P20GM121176). Lipidomics analysis was performed at Barshop Institute Functional Lipidomics Core funded by NIA, NIH (P30AG013319). The funding support was from NIGMS, NIH (T32GM144834 and R35GM147419).

## SUPPLEMENTAL INFORMATION

Supplemental Information includes 8 figures and 3 tables.

## Notes

### Competing Interest Statement

The authors have declared no competing interest.

### Summary of Updates

Changed title, added authors/changed author order, revised abstract, and included new results on the mechanism of ORF3a reducing BMP levels.

