## Supplementary material for "SARS-CoV-2 ORF3a blocks lysosomal cholesterol egress by disrupting VPS39-regulated NPC2 trafficking and BMP metabolism": Suppl 8 figures 3 tables

SUPPLEMENTAL INFORMATION

### Supplemental figures

**Figure S1**

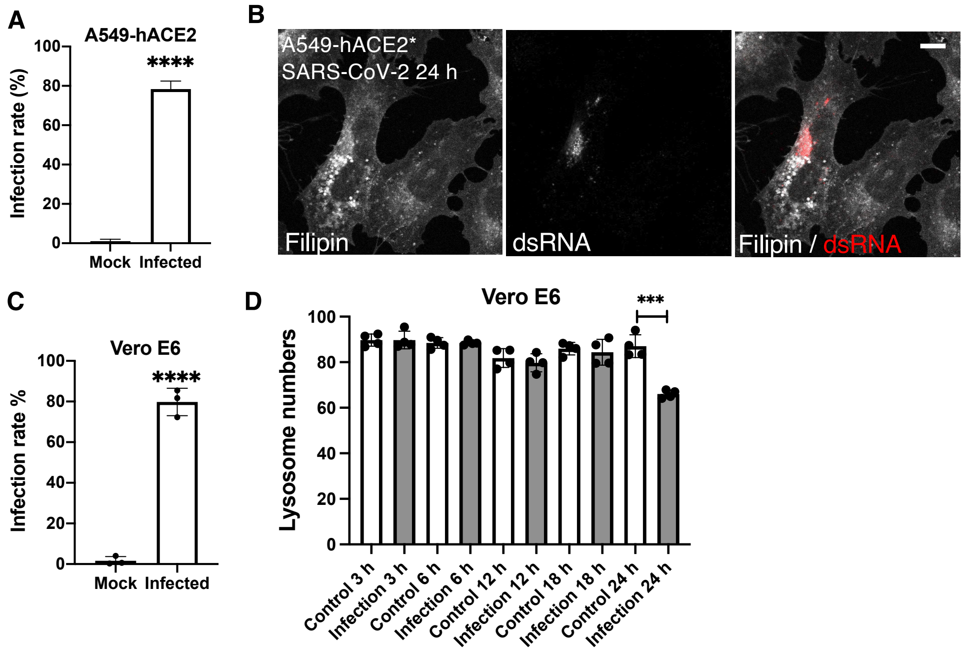


**Fig. S1. SARS-CoV-2 infection sequestrates cholesterol in lysosomes**

**A.** A549-hACE2 cells were infected with SARS-CoV-2, and the infection rate was determined by immunostaining of dsRNA and quantified using a high-content imaging system. **B.** An additional line of A549 cells stably expressing human ACE2 (A549-hACE2*) were infected with SARS-CoV-2, fixed at 24 h post-infection, stained with the antibodies against dsRNA and filipin, and imaged with a confocal microscope. Note that the dsRNA-positive cell showed increased filipin intensity, compared to the dsRNA-negative cell. Scale bars, 5 μm. **C.** Vero E6 cells were infected with SARS-CoV-2, and the infection rate was determined by immunostaining of dsRNA and quantified by high-content imaging system. D. Vero E6 cells were infected with SARS-CoV-2, fixed, immunostained with a LAMP1 antibody, and analyzed by high-content imaging. Bar graphs are presented as mean ± SD. $p$ values were determined using $t$ test. ***, $p$<0.001. ****, $p$<0.0001.

**Figure S2**

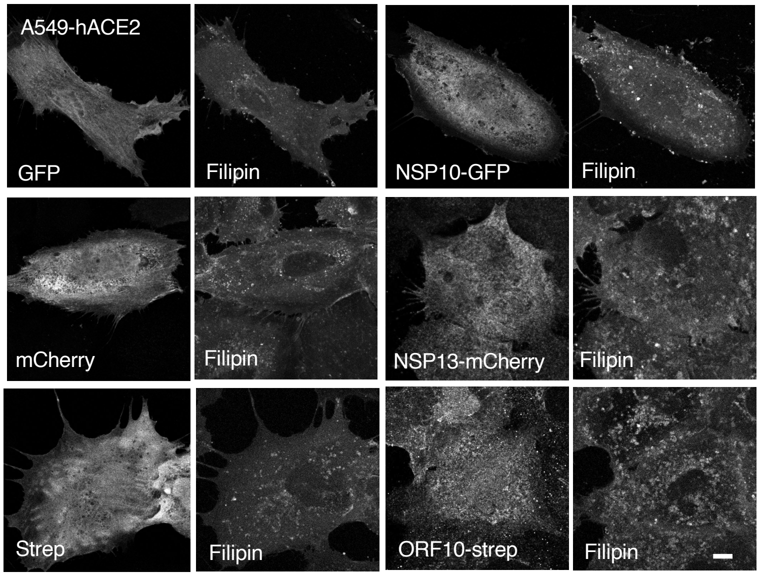


**Fig. S2. Cellular localization of NSP10, NSP13, and ORF10**

A549-hACE2 cells were transfected with the plasmids that encode NSP10, NSP13, or ORF10 with a different tag. Cells were fixed at 24 h post-transfection, stained with filipin and the antibodies against GFP, mCherry, or strep, and imaged with a confocal microscope. Scale bars, 10 μm.

**Figure S3**

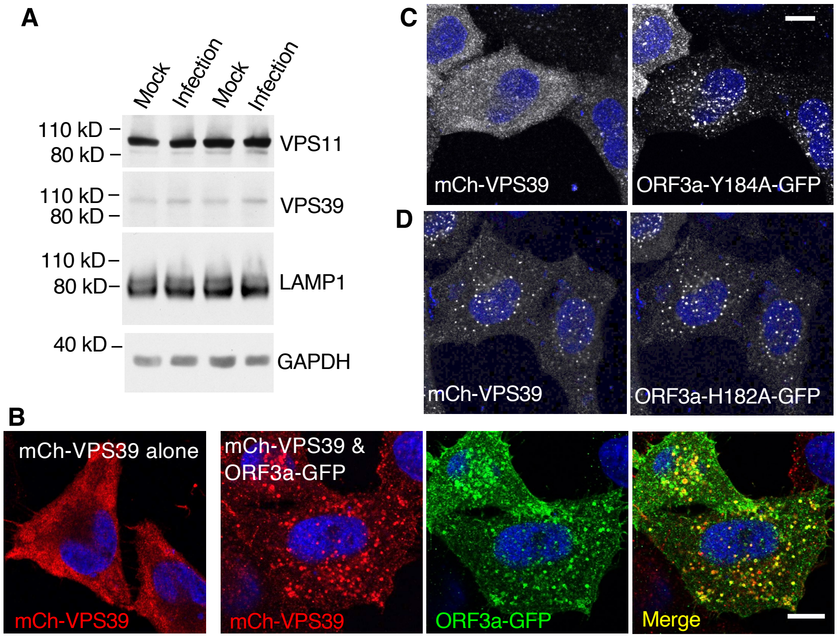


**Fig. S3. Characterization of ORF3a-VPS39 interaction**

**A.** A549-hACE2 cells were infected with SARS-CoV-2 and lysed at 24 h post-infection. Cell lysates were subjected to immunoblotting with the indicated antibodies. **B-D.** HeLa cells were transfected with mCh-VPS39 construct alone (**B, left**) or co-transfected with ORF3a-GFP (**B, right**) or ORF3a mutant constructs (**C,D**). The cells were fixed at 24 h post transfection, immunostained with the antibodies against GFP and mCherry, and imaged with a confocal microscope. Note that mCh-VPS39 alone displayed cytosolic distribution (**B, left**). When ORF3a was present, mCh-VPS39 formed puncta, colocalized with ORF3a (**B, right**). This distribution alteration was used as an indicator of ORF3a-VPS39 interaction in examining those ORF3a mutants (**C,D**). Scale bars, 5 µm.

**Figure S4**

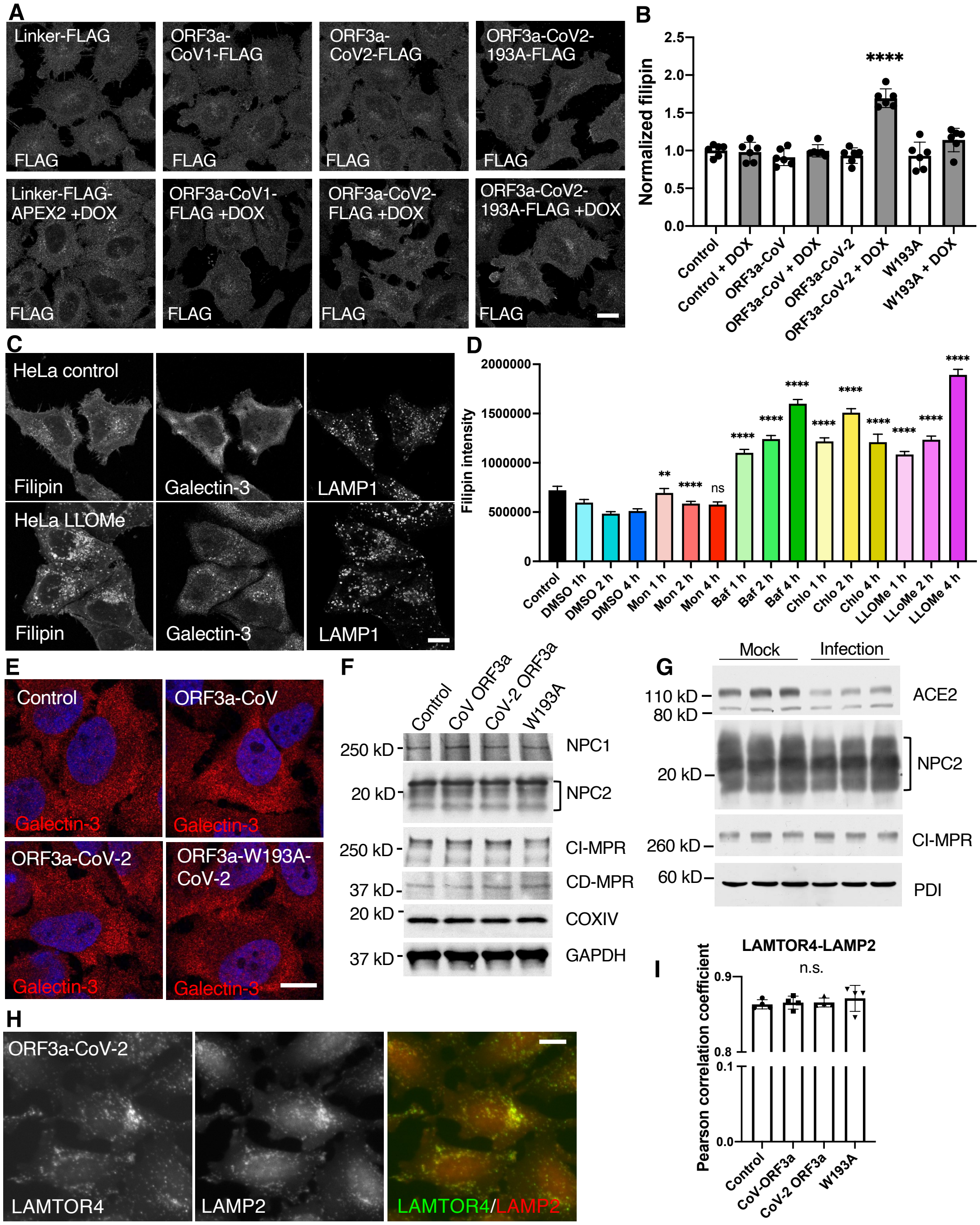


**Fig. S4. The impacts of ORF3a-VPS39 interaction on lysosome integrity**

**A.** HeLa Flp-In cells were treated with or without doxycycline (DOX), fixed, stained with a FLAG antibody, and imaged with a confocal microscope. **B**. The Flp-In cells were treated with or without doxycycline, fixed, stained with filipin, and analyzed by high-content imaging for filipin signals. **C.** HeLa cells were treated with 0.5 mM LLOMe for 1 h, fixed, stained with filipin and a galectin-3 antibody, and imaged with a confocal microscope. **D.** HeLa cells were treated with 10 $\mu$M monensin, 100 nM bafilomycin, 100 $\mu$M chloroquine, or 0.5 mM LLOMe for the indicated time periods, fixed, stained with filipin, and analyzed with high-content imaging. 0.1% DMSO in cell culture media served as a vehicle control. **E.** Example confocal microscopy images of HeLa Flp-In cells, immunostained with a galectin-3 antibody. Cell lysates from HeLa Flp-In cells (**F**) or SARS-CoV-2-infected A549-hACE2 cells (24 h post-infection, **G**) were subjected to immunoblotting with the indicated antibodies. **H,I.** HeLa Flp-In cells were immunostained with LAMTOR4 and LAMP2 antibodies and analyzed by high-content imaging. The colocalization between LAMTOR4 and LAMP2 was quantified and presented as Pearson correlation coefficient. Bar graphs are presented as mean ± SD. $p$ values were determined using one-way ANOVA. **, $p < 0.01$. ****, $p < 0.0001$. n.s., no significant difference. Scale bars, 5 μm.

**Figure S5**

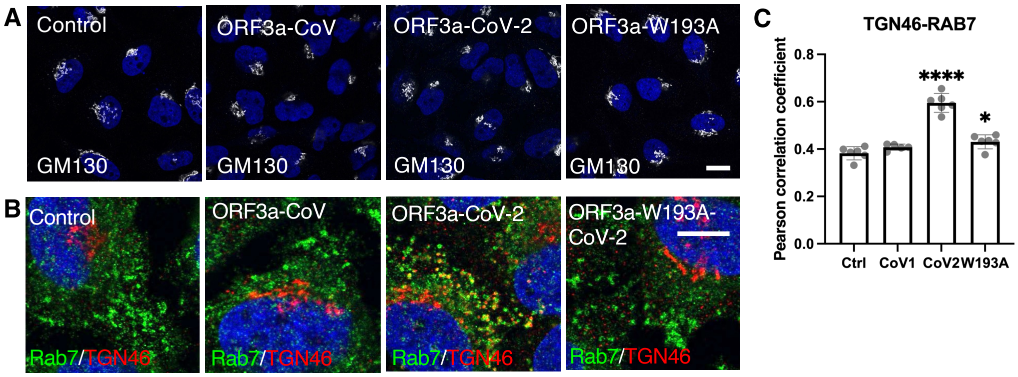


**Fig. S5. Characterization of Trans-Golgi Network (TGN) proteins**

HeLa Flp-In cells were fixed, immunostained with the indicated antibodies. Images were taken either with a confocal microscope **(A,B)** or high-content imaging system for quantification of the colocalization between TGN46 and Rab7 **(C).** Scale bars, 5 µm. Bar graphs are presented as mean ± SD. *p* values were determined using one-way ANOVA test. ****, $p < 0.0001$.

Figure S6

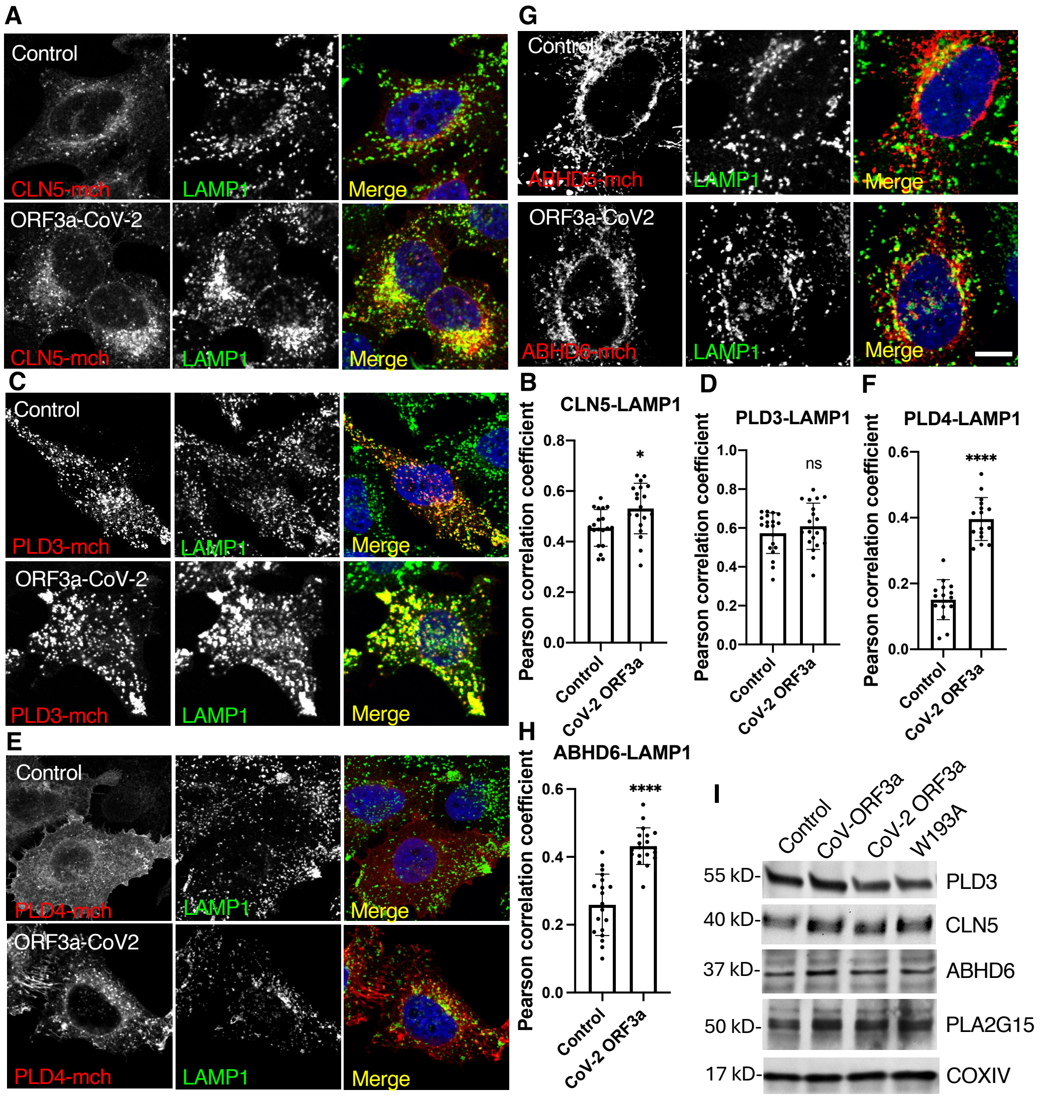


**Fig. S6. Characterization of BMP enzymes**

HeLa Flp-In control and CoV-2-ORF3a cells were transfected with the plasmids encoding BMP synthesis or turnover enzymes, fixed at 24-h post-transfection, immunostained with mCherry and LAMP1 antibodies. Images were taken with a confocal microscope **(A,C,E,G)** and used for quantification of the colocalization between the enzymes and LAMP1 by FIJI **(B,D,F,H).** Scale bar, 5 μm. **I.** Cell lysates from HeLa Flp-In cells were subjected to immunoblotting with the indicated antibodies. Bar graphs are presented as mean ± SD. $p$ values were determined using $t$ test. *, $p < 0.05$. ****, $p < 0.0001$. n.s., no significant difference.

**Figure S7**

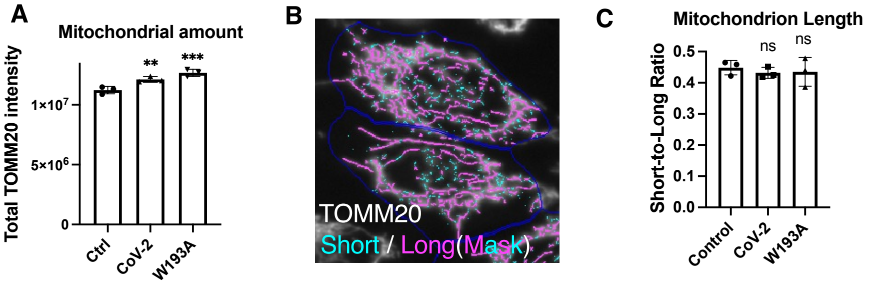


**Fig. S7. Mitochondrial quantifications**

HeLa Flp-In cells were fixed, immunostained with a TOMM20 antibody, and analyzed with high-content imaging. Average total intensity of TOMM20 (**A**) and the short-to-long ratio of mitochondria (**C**) were quantified. Short mitochondria were defined as those with a length shorter than 1 $\mu$m (**B**).

**Figure S8**

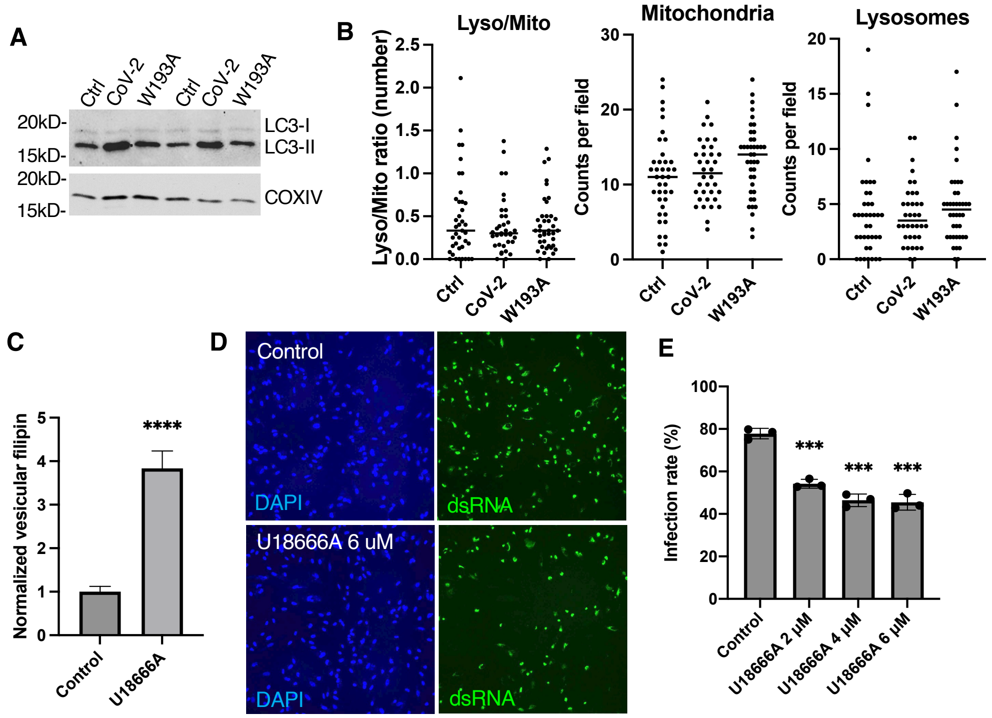


**Fig. S8. Quantification of lysosome-mitochondrion membrane contact sites and the effect of lysosomal cholesterol on SARS-CoV-2 infectivity**

**A.** Cell lysates from HeLa Flp-In cells were subjected to immunoblotting with the indicated antibodies. **B.** HeLa Flp-In cells were fixed and imaged by an electron microscope. The numbers of lysosomes and mitochondria were quantified from 40 images of 20 cells of each cell type. **C**. A549-hACE2 cells were treated with 2 $\mu$M U18666A for 4 h, fixed, stained with filipin, and quantified for filipin signals with high-content imaging. **D,E.** A549-hACE2 cells were treated with U18666A at the indicated concentration for 4 h and infected with SARS-CoV-2. Control cells were treated with the same amount of DMSO as in 6 $\mu$M U18666A for 4 h and infected equally. Cells were fixed at 24 h post-infection and immunostained with a dsRNA antibody to identify the infected cells. DAPI-stained nuclei were used to identify total cell numbers.

**Supplemental tables**

**Table. S1 cDNA constructs used in this study**

| **Insert** | **Vector** | **Tag** | **Remarks** |
|---|---|---|---|
| ABHD6 | pmCherry-N1 | c-mCherry | This study |
| CLN5 | pmCherry-N1 | c-mCherry | This study |
| GFP-E | pEGFPC1 | n-GPF | Addgne # 165123 |
| GFP-M | pEGFPC1 | n-GFP | Addgene #165124 |
| Linker peptide | pDEST-Flp-In | c-FLAG-APEX2 | This study |
| NPC2 | mcherry-N1 | c-mCherry | Gift from Dr. P. Lobel (Rutgers University) |
| SARS-CoV2-NSP1-strep | pLVX-EF1alpha-IRES-Puro | c-Strep | Addgene #141367 |
| NSP2-strep | pLVX-EF1alpha-IRES-Puro | c-Strep | Addgene #141368 |
| NSP3-GFP | pEGFP-N1 | c- GFP | Addgene #165108 |
| NSP4-mch | pmCherry-N1 | c-mcherry | Addgene #165132 |
| NSP5-strep | pLVX-EF1alpha-IRES-Puro | c-Strep | Addgene #141370 |
| NSP6-mch | pmCherry-N1 | c-mcherry | Addgene #165133 |
| GFP- NSP7 | pEGFP-C1 | n-GFP | Addgene #165112 |
| NSP8-GFP | pEGFP-N1 | c-GFP | Addgene #165113 |
| NSP9-strep | pLVX-EF1alpha-IRES-Puro | c-strep | Addgene #141375 |
| NSP10-GFP | pEGFP-N1 | c-GFP | Addgene #165115 |
| NSP11-strep | pLVX-EF1alpha-IRES-Puro | c-Strep | Addgene #141377 |
| GFP-NSP12 | pEGFP-C1 | n-GFP | Addgene #165117 |
| NSP13-mch | pmCherry-N1 | c-mcherry | Addgene #165136 |
| NSP14-mch | pmCherry-N1 | c-mcherry | Addgene #165137 |
| NSP15-GFP | pEGFP-C1 | n-GFP | Addgene #165120 |
| ORF3a-CoV-2-GFP | pEGFP-N1 | c-EGFP | This study |
| ORF3a-CoV-2-V5 | | | This study |
| CoV-ORF3a | pDEST-Flp-In | c-FLAG-APEX2 | This study |
| CoV-2-ORF3a-S171A | pEGFP-N1 | c-EGFP | This study |
| CoV-2-ORF3a-Y184A | pEGFP-N1 | c-EGFP | This study |
| CoV-2-ORF3a-H182A | pEGFP-N1 | c-EGFP | This study |
| CoV-2-ORF3a | pDEST-Flp-In | c-FLAG-APEX2 | Previous study |
| CoV-2-ORF3a-W193A | pDEST-Flp-In | c-FLAG-APEX2 | This study |
| CoV-2-ORF3a-W193K | pEGFP-N1 | c-EGFP | This study |
| CoV-2-ORF3a-W193H | pEGFP-N1 | c-EGFP | This study |

| | | | |
|---|---|---|---|
| CoV-2-ORF3a-W193D | pEGFP-N1 | c-EGFP | This study |
| CoV-2-ORF3a-W193F | pEGFP-N1 | c-EGFP | This study |
| ORF3a-CoV-GFP | pEGFP-N1 | c-GFP | Addgene # 165121 |
| ORF3b-strep | pLVX-EF1alpha-IRES-Puro | n-strep | Addgene #141384 |
| ORF6-strep | pLVX-EF1alpha-IRES-Puro | c-strep | Addgene #141387 |
| ORF7a-strep | pLVX-EF1alpha-IRES-Puro | c-strep | Addgene #141388 |
| ORF7b-strep | pLVX-EF1alpha-IRES-Puro | c-strep | Addgene #141389 |
| ORF8-GFP | pmCellFree_KA1 | n-GFP | Addgene 169395 |
| GFP-ORF9b | pEGFP-C1 | n-GFP | Addgene #165122 |
| ORF10-strep | pLVX-EF1alpha-IRES-Puro | c-Strep | Addgene #141394 |
| PLD3 | pmCherry-N1 | c-mCherry | This study |
| PLD4 | pmCherry-N1 | c-mCherry | This study |
| Spike | pGBW-m4137386 | No tag | Addgene 149540 |
| N-strep | pLVX-EF1alpha-IRES-Puro | c-Strep | Addgene #141391 |
| VPS39 | pmCherry-C1 | n-mcherry | Previous study |

**Table. S2 Antibodies used in this study**

| **Antibody** | **Source** | **Product Number** | **Dilution** | **Applications** |
|---|---|---|---|---|
| ABHD6 | Protein Tech | 20494-1-AP | 1:1000 | WB |
| ACE2 | Cell signaling technologies | 4355 | 1:1000 | WB |
| ATG5 | Protein Tech | 10181-2-AP | 1:1000 | WB |
| ATG7 | ThermoFisher Scientific | MA5-32221 | 1:1000 | WB |
| Calnexin | Cell Signaling Technologies | 2679s | 1:8000 | WB |
| Cathepsin D | Protein Tech | 219361 | 1:4000 | WB |
| CD-MPR | Developmental Studies Hybridoma Bank | 22d4 | 1:50 | WB |
| CI-MPR | Abcam | 124767 | 1:500 | IF |
| CI-MPR | Protein Tech | 20253-1-AP | 1:1000/1:500 | WB/IF |
| CLN5 | Abcam | ab170899 | 1:1000 | WB |
| COXIV | Cell Signaling Technologies | 11967S | 1:1000 | WB |
| dsRNA | Millipore Sigma | MABE1134 | | IF |
| DRP1 | Cell Signaling Technologies | 5391S | 1:1000 | WB |
| FLAG | Cell Signaling Technologies | 8146T | 1:1000 | WB |

| | | | | |
|---|---|---|---|---|
| FLAG (M2) | Millipore Sigma | A8592 | 1:1000 | IF |
| GAPDH | Protein Tech | 6004-1 | 1:5000 | WB |
| Galectin-3 | Biolegend | 125402 | 1:500 | IF |
| GFP-HRP | Miltenyi Biotec | 130-091-833 | 1:5000 | WB |
| GFP | ThermoFisher Scientific | A10262 | 1:1000 | IF |
| GM130 | Cell signaling technologies | 70767T | 1:500 | IF |
| LAMP1 | Cell Signaling Technologies | 9091s | 1:500 | IF |
| LAMP2 | Santa Cruz | SC-18822 | 1:500 | IF |
| LAMTOR4 | Cell Signaling Technologies | 13140S | 1:200 | IF |
| LBPA/BMP | Millipore Sigma | MABT837 | 1:500 | IF |
| LC3 | Santa Cruz | sc-271625 | 1:1000 | WB |
| mCherry | ThermoFisher Scientific | M11217 | 1:1000/1:5,000 | IF/WB |
| MIRO1 | Millipore Sigma | HPA010687 | 1:1000 | WB |
| MIRO2 | Protein Tech | 11237-1-AP | 1:1000 | WB |
| NPC1 | Abcam | 134113 | 1:1000/1:500 | WB/IF |
| NPC2 | Gift from P. Lobel | | 1:10000 | WB |
| PDI | Cell Signaling Technologies | 3501S | 1:10,000 | WB |
| PDH | Abcam | ab110333 | 1:500 | IF |
| PLA2G15 (LYPLA3) | Santa Cruz | Sc-376078 | 1:500 | WB |
| Rab7a | Cell Signaling Technologies | 9367 | 1:200 | IF |
| STX17 | Abcam | ab316119 | 1:1000 | WB |
| TGN46 | BioRad | AHP500GT | 1:500 | IF |
| TOMM20 | Santa Cruz | sc-17764 | 1:500 | IF |
| TOMM20 | Cell Signaling Technologies | 42406 | 1:2000 | WB |
| ULK1 | Cell Signaling Technologies | 8054S | 1:1000 | WB |
| V5 | Protein Tech | 14440-1-AP | 1:500 | IF |
| VPS11 | Santa Cruz | sc-515094 | 1:1000 | WB |
| VPS29 | Cell Signaling Technologies | 73540 | 1:1000 | WB |
| VPS35 | Abcam | Ab10099 | 1:1000/1;500 | WB/IF |
| VPS39 | Santa Cruz | SC-514762 | 1:500 | WB |
| VPS41 | Santa Cruz | SC-377118 | 1:500 | WB |

**Table. S3 Other important reagents used in this study**

| Reagent | Source | Product Number | Applications |
|---|---|---|---|
| Bafilomycin | Milipore Sigma | 88899-55-2 | V-ATPase inhibitor |

| | | | |
|---|---|---|---|
| (S,S) LBPA/BMP | Echelon bioscience | L-B181 | BMP addition |
| CellMask | Thermos fisher | H32722 | Labels entire cells |
| Chloroquine | Milipore Sigma | 50-63-5 | V-ATPase inhibitor |
| Filipin | Milipore Sigma | F9765 | Probes free cholesterol |
| LLOMe | Milipore Sigma | 16689-14-8 | Lysosome damage reagent |
| Monencin | Milipore Sigma | 1445481 | Increases lysosome pH |
| siMIRO1 | Horizon discovery | L-010365-01-0005 | RNAi |
| siMIRO2 | Horizon discovery | L-008340-01-0005 | RNAi |
| siDRP1 | Horizon discovery | L-012092-00-0005 | RNAi |
| siATG5 | Horizon discovery | J-004374-07-0002 | RNAi |
| siATG7 | Horizon discovery | L-020112-00-0005 | RNAi |
| siULK1 | Horizon discovery | J-005049-05-0002 | RNAi |
| U18666A | Milipore Sigma | U3633 | NPC1 inhibitor |
